# Hybridization outcomes reflect context-dependent reproductive isolation in two damselfly hybrid zones

**DOI:** 10.64898/2025.12.23.696213

**Authors:** Luis Rodrigo Arce-Valdés, Janne Swaegers, Andrea Ballén-Guapacha, Jesús Ramsés Chávez-Ríos, Pallavi Chauhan, Maren Wellenreuther, Bengt Hansson, Rosa Ana Sánchez-Guillén

## Abstract

Pair of species with multiple hybrid zones provide powerful systems to examine how reproductive isolation and reinforcement may depend on ecological and demographic contexts. We conducted a comparative analysis of two hybrid zones between *Ischnura elegans* and *Ischnura graellsii* in North-central (NC) and North-western (NW) Spain to evaluate how differences in reproductive barriers shape patterns of hybridization and admixture. First, we quantified five prezygotic barriers in the NC hybrid zone—where no barrier estimates previously existed—and compared them with published data from the NW hybrid zone. Second, we conducted genomic analyses using 5,702 SNPs to assess genetic structure, differentiation, and diversity between species; and 381 SNPs with species-specific alleles to characterize hybrid classes and introgression across both regions. Reproductive isolation was strongly context dependent. Mechanical reinforcement acted in the same cross direction in both hybrid zones, but the NC hybrid zone also showed stronger gametic (fertility) isolation. Consequently, total prezygotic isolation was higher in the NC hybrid zone. These differences were also observed genomically: the NC hybrid zone exhibited limited and largely unidirectional introgression, whereas the NW hybrid zone showed more extensive admixture, bidirectional gene flow, and reduced interspecific differentiation, consistent with a longer or more sustained history of hybridization. Our results demonstrate that hybridization outcomes can differ markedly between hybrid zones involving the same pair of species, underscoring the importance of evaluating spatial variation in reproductive barriers and introgression to assess the presence, asymmetry, and repeatability of reinforcement, as well as the microevolutionary processes shaping hybrid zone structure.

## INTRODUCTION

Hybridization between closely related species can lead to a wide range of evolutionary outcomes, including interspecific gene flow, reinforcement of reproductive barriers, species fusion or even the origin of a new species (Seehausen, 2004). Among these outcomes reinforcement is a key mechanism promoting reproductive isolation. It occurs when natural selection against low-fitness hybrids strengthens prezygotic barriers upon secondary contact (Dobzhansky, 1937; Noor, 1999; Coyne and Orr, 2004; Lukhtanov, 2011). This often leads to Reproductive Character Displacement (RCD), i.e., greater divergence in reproductive traits in sympatry than in allopatry (Howard, 1993). Reinforcement can also be asymmetric, affecting primarily one reciprocal cross, one species or one sex, depending on the costs of hybridization and the traits involved (Servedio and Noor, 2003; Turelli and Moyle, 2007; Yukilevich, 2012).

The nature of the outcomes of hybridization depends on both intrinsic and extrinsic factors (Seehausen, 2004; Abbott *et al*., 2013; Todesco *et al*., 2016). Intrinsic factors include species-specific traits such as the degree of reproductive isolation and genetic architecture, while extrinsic factors include ecological and demographic conditions such as habitat overlap, population densities, and environmentally dependent hybrid fitness (Liou and Price, 1994; Coyne and Orr, 2004; Yukilevich, 2012; Walsh *et al*., 2016). Empirical studies reveal that the strength and asymmetry of the reproductive barriers, as well as patterns of introgression, often differ across independent hybrid zones, reflecting both intrinsic traits and environmental conditions (Buerkle and Rieseberg, 2001; Vines *et al*., 2003; Sweigart *et al*., 2007; Good *et al*., 2008; Nolte *et al*., 2009; Aboim *et al*., 2010; Haselhorst and Buerkle, 2013; Gompert *et al*., 2014; Mandeville *et al*., 2017). This context-dependence has been documented across plants (e.g. Haselhorst and Buerkle, 2013), insects (e.g. Larson *et al*., 2014), fishes (e.g. Aboim *et al*., 2010), amphibians (e.g. Vines *et al*., 2003), and mammals (e.g. Chavez *et al*., 2011), highlighting that hybridization outcomes can differ dramatically and rarely show complete consistency among zones. Nonetheless, some comparative studies—such as Buerkle and Rieseberg (2001) in sunflowers (*Helianthus annuus* and *H. petiolaris*)—report remarkably consistent patterns of introgression, suggesting that intrinsic genetic factors can predominate over local environmental effects. Therefore, examining multiple hybrid zones involving the same pair of species offers a powerful approach to test the repeatability of hybridization dynamics and the presence of reinforcement (Abbott *et al*., 2013; Harrison and Larson, 2016).

In insects, ongoing climate change and anthropogenic pressures are driving shifts in species distributions creating new zones of sympatry or altering existing ones (Sánchez-Guillén *et al*., 2016; González-Tokman *et al*., 2020; Arce-Valdés and Sánchez-Guillén, 2022). These changes modify the conditions under which species interact, providing natural experiments to investigate how hybridization outcomes vary across ecological and demographic contexts. Despite its importance, the repeatability and context-dependence of hybridization outcomes remain poorly understood, particularly in animal systems. Odonates represent an ideal system to explore these questions, as they are heavily sensitive to rising temperatures, with many species shifting their distributions in response to climate change (Hickling *et al*., 2005; Hassall *et al*., 2007; Hassall and Thompson, 2008; Ott, 2010; Lancaster *et al*., 2016; Sánchez-Guillén *et al*., 2023).The damselfly *Ischnura elegans* has expanded its range both northwards (into Britain, Cham *et al*., 2014; and into Sweden, Dudaniec *et al*., 2018) and southwards (into Spain, Sánchez-Guillén *et al*., 2011). This southward expansion has brought *I. elegans* into increasing contact with *Ischnura graellsii,* a closely related species with which it shares numerous genetic, phenotypic, and ecological traits (Monetti *et al*., 2002; Sánchez-Guillén *et al*., 2011; Wellenreuther *et al*., 2018). *Ischnura graellsii* is generally very abundant throughout the Iberian Peninsula, whereas *I. elegans* has a fragmented and patchy distribution (Ocharan-Larrondo, 1987). This expansion has been associated with the adaptation of *I. elegans* to the Spanish thermal regime, a process likely facilitated by phenotypic plasticity and epigenetic mechanisms (Swaegers, Sánchez-Guillén, Carbonell, *et al*., 2022; Swaegers *et al*., 2024). Consequently, *I. elegans* has undergone an environmental niche shift, bringing it into closer alignment with the niche already occupied by *I. graellsii* (Wellenreuther *et al*., 2018). This increasing overlap between the two species often results in the replacement of *I. graellsii* by *I. elegans*, particularly along the Mediterranean coast (Ocharan-Larrondo, 1987; Sánchez-Guillén *et al*., 2011). As a consequence of this pronounced demographic displacement, *I. graellsii* has recently been classified as a vulnerable species (De Knijf *et al*., 2024).

As a result of this secondary contact, *I. elegans* and *I. graellsii* form hybrid zones in several regions of Spain. This provides a natural context to study heterospecific reproductive interactions and the mechanisms limiting gene flow. In the North-west (NW) hybrid zone, reproductive isolation between the two species is highly asymmetric between reciprocal crosses, resulting from the joint action of mechanical barriers, gametic incompatibilities, and hybrid incompatibilities (Sánchez-Guillén *et al*., 2012). This asymmetry arises primarily from reinforcement acting asymmetrically on the mechanical barrier involved in the tandem formation —the initial contact during mating involving secondary genitalia— in the cross between *I. graellsii* males and *I. elegans* females (Arce-Valdés *et al*., 2024). Thus, reinforcement limits gene flow in only one direction, while allowing hybridization to persist in the reciprocal cross. Consistent with this, reproductive character displacement has been detected exclusively in tandem-related traits of *I. graellsii* males and *I. elegans* females (Ballén-Guapacha *et al*., 2024), suggesting that selection is acting specifically on this cross to reduce maladaptive hybridization. This pattern is expected under asymmetric reinforcement (Servedio and Noor, 2003). Although a similar pattern of reproductive character displacement has been detected in *I. elegans* females from the north-central (NC) hybrid zone (see Ballen-Guapacha *et al*., 2023), reproductive isolation and reinforcement have not yet been formally assessed in this region. At the genetic level, early studies based on a limited set of microsatellite markers focused on introgression into *I. elegans* and reported high levels of admixture and introgression (Sánchez-Guillén *et al*., 2011; Wellenreuther *et al*., 2018), although such estimates may have been inflated due to molecular markers limitations (see Miralles *et al*., 2023). In contrast, a more recent genomic-wide SNP study (Swaegers, Sánchez-Guillén, Chauhan, *et al*., 2022) examined patterns of restricted introgression on sex chromosomes but did not explore broader dynamics of gene flow or reproductive barriers. Consequently, key questions remain open in both hybrid zones —particularly in the NC hybrid zone— including the directionality and extent of introgression, the strength and symmetry of reproductive isolation, and the role of reinforcement.

In this study, we examined two hybrid zones between *I. elegans* and *I. graellsii*, which differ in both the timing and historical context of secondary contact: the NC hybrid zone reflects a recent contact (∼36–54 generations ago, ∼18 years) in a region still connected to allopatric populations, whereas the NW hybrid zone represents an older contact (∼60–80 generations ago, ∼30 years) that occurred in an isolated region (Sánchez-Guillén *et al*., 2011). By integrating experimental crosses and genome-wide SNP data, our goal was to evaluate whether variation in the strength of reproductive barriers across these hybrid zones aligns with differences in the direction and extent of gene flow, interspecific differentiation, genetic diversity as well as the repeatability of reinforcement—specifically, whether reinforcement occurs, and is asymmetric, in the NC hybrid zone. To this end, (i) we quantified five reproductive barriers between *I. elegans* and *I. graellsii* in the NC hybrid zone—where no prior data existed, unlike the NW zone—and (ii) we analyzed genome-wide SNP data to characterize patterns of interspecific introgression, genetic structure, diversity, and differentiation in both hybrid zones. We hypothesized that both the differences in the timing of secondary contact(assuming that older hybrid zones provide more opportunity for the strengthening of prezygotic reproductive barriers through reinforcement and for introgression via interspecific gene flow) and in connectivity to allopatric populations, would shape differential evolution of reproductive isolation between hybrid zones. In particular, we expect the NC hybrid zone to show (i) reinforcement of the mechanical barrier involved in tandem formation—especially in crosses between *I. graellsii* males and *I. elegans* females—, (ii) higher levels of postmating isolation compared to the NW hybrid zone, (iii) a reduced prevalence of hybrid and introgressed individuals, particularly in the cross direction where reinforcement occurs due to stronger reproductive isolation, (iv) lower intraspecific genetic structure and differentiation than the NW hybrid zone, but (v) higher interspecific genetic differentiation and (vi) lower levels of genetic diversity than the NW hybrid zone due to its shorter history of hybridization and introgression.

## MATERIAL AND METHODS

### Reproductive isolation in the NC hybrid zone

#### Reproduction in Ischnura

Reproduction in *Ischnura* damselflies involves a male actively searching for a female, and in some species also courting her, a process driven by sexual selection (Fincke, 1997). Once the male locates a potential mate, he grasps the female by her prothorax using his caudal appendages (tandem position). This leads to copulation if the female accepts the male by bending her abdomen towards him, allowing contact between the male and female genitals (wheel position; Corbet, 1999). During copulation, three behavioral phases defined by internal genital activity follow (Miller and Miller, 1981). First, males remove sperm from previous matings from the bursa and spermatheca (Miller, 1987a, 1987b; Cordero and Miller, 1992), copulation progresses through insemination, followed by male mate guarding, terminating insemination (Cordero-Rivera *et al*., 2010). After copulation, the female lay eggs until her sperm reserves are depleted or until she engages in subsequent matings.

#### Prezygotic barriers in the NC hybrid zone

Last-instar larvae of approximately 200 *I. elegans* and 200 *I. graellsii* individuals collected from the NC hybrid zone (Mateo, Valbornedo and Villar), were sampled in June-July of 2016, 2017, and 2018. Larvae were transported to the laboratory and maintained until adulthood (for details about larval rearing methodology see Van Gossum *et al*., 2003; Sánchez-Guillén *et al*., 2005). Species identity was confirmed upon emergence at the adult stage based on morphological traits: male caudal appendages, thorax color in young females, and the shape of the prothoracic tubercle in both sexes (see Monetti *et al*., 2002). Individuals from each locality were kept isolated, and detailed records of their origin were maintained throughout the experiments. Crosses were performed using individuals from all three localities in all possible combinations (e.g., Mateo × Valbornedo, Mateo × Villar, Villar × Valbornedo), both within and between species. For heterospecific crosses, we included both reciprocal directions: *I. elegans* males × *I. graellsii* females and *I. graellsii* males × *I. elegans* females. Similar data from allopatry and the NW hybrid zone are available from two previous studies (Sánchez-Guillén *et al*., 2012; Arce-Valdés *et al*., 2024).

We measured five prezygotic barriers—two premating, mechanical (tandem position) and mechanical-tactile (wheel position), and three postmating (oviposition, fecundity and fertility) barriers. To assess (1) the mechanical barrier we measured the incompatibility between the male cerci and the female prothorax, indicating the male’s inability to grasp the female into a tandem position. Then, if a successful tandem was formed, we measured (2) the mechanical-tactile barrier as the frequency of tandems that progressed to copulation (wheel position). This assessed female rejection of copulation or the incompatibility between male and female primary genitalia. We evaluated oviposition, fecundity, and fertility to assess various factors impeding fertilization post-copulation, such as inefficient sperm transfer or storage, sperm incapacity in foreign reproductive tracts, gamete failure to initiate fertilization upon contact, and the foreign ejaculate’s inability to induce or reduce oviposition rates (see Coyne and Orr, 2004). (3) Oviposition success was assessed by comparing the percentage of *I. elegans* and *I. graellsii* females that laid eggs after conspecific and heterospecific matings, respectively. (4) Fecundity was determined by the average number of eggs laid across the first three clutches, while (5) fertility was quantified as the average number of fertile eggs in each mating treatment, considering only eggs that hatched or showed embryo development. See Arce-Valdés *et al*. (2024) for additional details on reproductive barriers measurements in *Ischnura*.

#### Statistical comparisons of prezygotic isolation between hybrid zones

Our primary objective was to compare the probability of gene flow between *I. elegans* and *I. graellsii* among allopatry and the two hybrid zones (NW and NC). For the NC hybrid zone, we used empirical data generated in the present study. For the NW hybrid zone and the allopatric populations, we relied on previously published data from Sánchez-Guillén *et al*. (2012) and Arce-Valdés *et al*. (2024).

To quantify the proportional decrease in hybridization relative to random mating, we applied to each reproductive barrier the formula proposed by Sobel and Chen (Sobel and Chen, 2014):

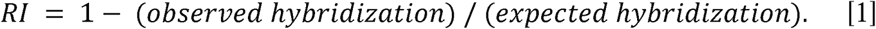

This formula facilitates the calculation of average values, variances, and confidence intervals, providing insights into the potential range of average reproductive isolation.

To estimate cumulative reproductive isolation between *I. graellsii* and *I. elegans*, we employed a multiplicative function that integrates the sequential effects of individual reproductive barriers, as described by Coyne and Orr (1989, 1997) and Ramsey *et al*. (2003). The cumulative contribution (CC) of a component to reproductive isolation (RI) at stage n was calculated as follows:

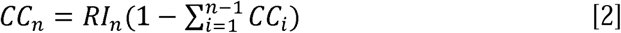

To evaluate the effects of different cross types (♂E×♀G, ♂G×♀E) and zones (allopatry, NW and NC) on RI we used generalized linear models (GLMs). GLMs were modeled using the glm() function in R 4.3.0 (R Core Team, 2024) and compared using AICc values with the dredge() function of the MuMIN 1.47.5 library (Barton, 2009). The mechanical (successful tandem = 1 *vs.* unsuccessful tandem = 0), mechanical-tactile (successful mating = 1 *vs.* unsuccessful mating = 0), oviposition (mated female that laid eggs = 1 *vs.* mated female that did not lay eggs = 0), and fertility (fertile egg = 1 *vs.* unfertile egg = 0) barriers were modeled using the binomial distribution, while the fecundity barrier (average number of laid eggs in the first three clutches) was modeled using the gamma distribution (Table 1). The most parsimonious model per reproductive barrier was selected based on the lowest AICc score. Goodness-of-fit of each selected model was assessed by simulating its residuals using the DHARMa 0.4.6 library (Fig. S1; Hartig and Lohse, 2022). Pairwise statistical comparisons were conducted for the variables included in the most probable model. Since we followed the same methods as Arce-Valdés *et al*. (2024), readers may refer to that manuscript for additional details.

**Table 1.** Summary of reproductive barriers GLMs and *post hoc* Tuckey pairwise test p values. Reproductive 994 isolation was measured following Arce-Valdés *et al*. (2024).

| Reproductive barrier | GLM response variable | Best GLM following AICc | ♂ *I. elegans* × ♀ *I. graellsii* |  |  | ♂ *I. graellsii* × ♀ *I. elegans* |  |  |
| --- | --- | --- | --- | --- | --- | --- | --- | --- |
|  |  |  | Allop. vs NW | Allop. vs NC | NW vs NC | Allop. vs NW | Allop. vs NC | NW vs NC |
| Mechanical | Binomial: number of successful tandems vs number of unsuccessful tandem attempts | RI ~ Crosses + Distribution + (Crosses * Distribution) | 0.8074 | 0.5292 | 0.8128 | **0.0002\*** | **0.0289\*** | 0.1303 |
| Mechanical-Tactile | Binomial: number of successful matings vs number of unsuccessful mating attempts | Null model | 0.9482 | 0.7057 | 0.8281 | 0.5125 | 0.9610 | 0.7070 |
| Oviposition | Binomial: number of mated females that laid eggs vs number of mated females that did not lay eggs | RI ~ Crosses + Distribution | **0.0025\*** | 0.5005 | 0.3941 | 1.0000 | 0.9999 | 1.0000 |
| Fecundity | Gamma: Average number of eggs laid during the first three clutches per female. | RI ~ Crosses + Distribution + (Crosses * Distribution) | **0.0178\*** | 0.4212 | 0.2171 | 0.8281 | 0.9995 | 0.8423 |
| Fertility | Binomial: number of fertile eggs vs number of infertile eggs. | RI ~ Crosses + Distribution + (Crosses * Distribution) | **<0.0001\*** | **<0.0001\*** | **<0.0001\*** | 0.9307 | **<0.0001\*** | **<0.0001\*** |
| * p < 0.05. |  |  |  |  |  |  |  |  |

### Genetic consequences of hybridization across hybrid zones

#### Population selection for genomic analyses

Genomic analyses were based on five single-digest RAD libraries provided by Swaegers *et al*. (2022). From the populations included in the five libraries generated by Swaegers *et al*. (2022), we selected a subset of populations sampled across the distribution of *I. elegans* and *I. graellsii*, with the aim of capturing both allopatric and sympatric distributions. From allopatry, we included five populations of *I. elegans* and three of *I. graellsii* (Table S1) to cover a broad latitudinal gradient in order to capture the genetic diversity of both species as full as possible. In sympatry, we included populations representing three scenarios: (1) dominance of *I. elegans*, (2) dominance of *I. graellsii*, and (3) mixed proportions of both species (Table S1). Specifically, from the NC hybrid zone (Table S1), we included seven populations (Arreo, Cañas, Mateo, Perdiguero, Valbornedo, Valpierre, and Villar), and from the NW hybrid zone (Table S1), we included four populations (Doniños, Laxe, Louro, and Xuño).

This sampling strategy was designed to ensure the detection of hybrids and backcrosses in both directions. Because hybridization and introgression are often biased towards the rare species—due to it having a higher probability for heterospecific encounters than the common species (Yukilevich, 2012)—we deliberately included populations where either species was dominant. This approach enabled us to evaluate not only the occurrence of hybridization but also the direction and extent of introgression. Individuals were initially assigned to either *I. elegans* or *I. graellsii*. While species identification in females is less reliable, we included them to avoid underestimating admixture, particularly in light of the Haldane’s rule (Swaegers, Sánchez-Guillén, Chauhan, *et al*., 2022). Additionally, we included a population from Menorca (Albufeira, Balearic Islands) to assess the replacement of *I. graellsii* by *I. elegans* in that region (Table S1).

#### SNP calling and quality filtering

Single digest RAD libraries were processed using the STACKS v2.2 pipeline (Catchen *et al*., 2013; Rochette *et al*., 2019). Raw reads were demultiplexed with *process_radtags* and PCR clones were identified and discarded with *clone_filter* using the default parameters. Sequence reads were aligned to the *I. elegans* draft genome assembly (Chauhan *et al*., 2021) using BOWTIE2 v.2.3 (mismatch allowance per seed alignment of 1, maximum mismatch penalty of 6 and minimum of 2, maximum fragment length of 1000 bp and minimum of 100 bp; Langmead and Salzberg, 2012). We used the *ref_map* pipeline to detect SNPs using default parameters. Only SNPs with a minor allele frequency of less than 0.05 and a maximum observed heterozygosity of 0.70 were retained. Moreover, the locus had to occur in 80% of the individuals in a population and in at least 18 of the 20 populations to be included in the final SNP dataset. SNP markers were filtered to include only a single SNP on each RAD tag to create a data set without closely linked loci (using the **--***write_single_snp* option in STACKS). Finally, using the *I. elegans* reference genome (Chauhan *et al*., 2021) SNPs were filtered to include only those located on autosomal scaffolds. Exploratory analyses of population structure revealed possible hybridization in two of the *I. graellsii* samples from Seyhouse (Algeria), probably with a third *Ischnura* species (Fig. S2). These two samples were removed from further analyses leaving a final total sample size of 185 (Table S1).

#### Identification of diagnostic SNPs with species-specific alleles

Markers with diagnostic species-specific alleles are useful for assigning later-generation hybrids and detecting introgressed alleles in population genetic studies (Hohenlohe *et al*., 2011). To provide a list of such markers, alternatively fixed SNPs between the allopatric populations of the parental species were identified using VCFtools v0.1.16 (Danecek *et al*., 2011). These analyses were based on 43 *I. elegans* and 25 *I. graellsii* individuals. SNPs for each allopatric zone that showed only one allele (*--max-maf* 0) were selected, and then, shared loci between the two allopatric zones were identified using the *intersect()* function of R (R Core Team, 2024). Next, we applied a Hardy-Weinberg equilibrium test to these loci using VCFtools (*--hardy*) and excluded those fixed for the same allele in both species (H_E_=0). The final dataset consisted of 381 SNPs with species-specific alleles (out of the 5,702 total SNPs), all with F_ST_=1. Note that this set of SNPs with species-specific alleles might not fully represent fixed alleles, but rather alleles with highly skewed frequencies between our species groups (Fitzpatrick, 2012; Jordan *et al*., 2018). We will refer henceforth to this dataset as “diagnostic SNPs” as it has been employed in the literature (Hohenlohe *et al*., 2011).

#### Genetic structure analyses

We used ADMIXTURE v1.3.0 (Alexander and Lange, 2011) to assess genetic structure based on two SNP datasets: the full set of 5,702 SNPs and the subset of 381 diagnostic SNPs. Analyses were first conducted under the “unsupervised model” which does not employ reference populations for ancestry assignment. For the full dataset, we evaluated K (number of ancestral genetic clusters) from 1 to 21 (corresponding to the number of sampled populations plus one) and identified the optimal *K* as the value with the lowest cross-validation (cv) error. The same unsupervised model was applied to the subset of diagnostic SNPs subset with K values ranging from 1 to 15. This reduced range reflects the grouping of allopatric samples into a single group fixed for alternative alleles per species. Finally, we ran ADMIXTURE under the “supervised model” using the allopatric populations of each species as our two reference-groups (K=2), for both the full and the diagnostic SNPs dataset, to further evaluate patterns of ancestry and population structure. We also visualized patterns of genetic structure using a Principal Component Analysis (PCA) with the R package SNPRelate v1.6.4, function *snpgdsPCA()* (Zheng *et al*., 2012), using the set of 5,702 SNPs.

#### Admixture and assignment to hybrid classes

We used the R package INTROGRESS v1.2.3 (Gompert and Buerkle, 2010) to calculate individual introgression coefficients, hybrid index (HI-values) and individual heterozygosity (HET-values), and used both to classify individuals into different hybrid classes (Jordan *et al*., 2018). INTROGRESS was used with the dataset of 381 diagnostic SNPs, as the assignment to hybrid classes can be inexact when using non-diagnostic markers (Buerkle, 2005). When using SNPs fixed for alternative alleles between parental species, INTROGRESS calculates the hybrid index as the proportion of alleles inherited from one species and the heterozygosity as the proportion of alleles that are heterozygous, ranging from 0 (pure species) to 1 (F_1_ hybrids). Individuals of pure species are expected to be 100% homozygous and F_1_ hybrids are 100% heterozygous (Gompert and Buerkle, 2010). Thus, the HI-value gives the proportion of alleles inherited from one species, in this case *I. elegans* (e.g., 1.00=100% *I. elegans*, and 0.00=100% *I. graellsii*, alleles), whereas HET-values, which range from 0.00 to 1.00 (0.00=all sites are homozygous, 1.00=all sites are heterozygous) are used as a proxy of the number of generations since a hybridization event occurred within the ancestry of each individual. First-generation hybrids (F_1_ individuals) are expected to be heterozygous at all species-specific alleles SNPs, while later-generation hybrids and backcrosses would have lower heterozygosity levels. Additionally, the HI-values of F_1_ and F_2_ individuals will be close to 0.5, while backcrosses will have a HI-value below or above 0.5 (Fitzpatrick, 2012).

However, due to model uncertainty (Mandeville *et al*., 2017) we, conservatively, considered individuals with HI < 5% as pure *I. graellsii* and those with HI > 95% as pure *I. elegans*. Individuals with HI between 5–10% were classified as introgressed *I. graellsii*, while those with HI between 90–95% were classified as introgressed *I. elegans*. For consistency, we also relaxed the criteria for F_1_ and F_2_ hybrid classes; we classified individuals into eight parental and hybrid classes (cf. Milne and Abbott, 2008; Walsh *et al*., 2015): (i) pure *I. elegans* (HI=0.95-1.000; HET≤0.08), (ii) pure *I. graellsii* (HI=0.000-0.05; HET≤0.08), (iii) introgressed-*elegans* (HI=0.900-0.950; HET≤0.16), (iv) introgressed-*graellsii* (HI=0.05-0.100; HET≤0.16), (v) backcross-*elegans* (HI=0.601-0.899; HET=0.118-0.449), (vi) backcross-*graellsii* (HI=0.101-0.399; HET=0.118-0.449), (vii) relaxed F_1_ hybrids (HI=0.400-0.600; HET≥0.700), and (viii) relaxed F_2_ hybrids (HI=0.400-0.600; HET=0.450-0.69).

#### Testing for deviations from neutral hybrid class expectations

To investigate whether the NW and NC hybrid zones differ in their hybrid class composition, we compared the observed distributions to a neutral expectation where all hybrid classes are assumed to be equally common. This null model represents a scenario without selection or reproductive barriers. Expected frequencies were calculated using a contingency table, and the observed admixture-class distribution in both zones was compared to this prediction. Z-tests with Yates’s correction for small sample sizes were used to asses differences in the proportions of each hybrid class category.

#### Qualitative characterization of hybrid zone interspecific introgression

To better interpret the biological significance of these differences in hybrid class composition, we further categorized the populations from both hybrid zones into three qualitative hybridization patterns based on their genotypic profiles. This classification was determined by the frequency distribution of the different hybrid classes: 1) introgressed hybridization pattern, where the distribution ranges from introgressed to pure individuals, with the mode strongly skewed towards one of the parental species; 2) unimodal hybridization pattern, where the distribution includes a wide range of hybrid, admixed, and backcrossed genotypes, often toward one or both parental species; and, 3) bimodal hybridization pattern, where the distribution is bimodal, dominated by the two parental genotypes with few hybrids (F_1_ and F_2_ hybrids) present (Jiggins and Mallet, 2000).

#### Genetic diversity across hybrid zones

We examined whether hybridization has contributed to an increment in the genetic diversity in sympatry by comparing estimates across allopatric and sympatric populations of each species. For each locality, we calculated diversity metrics both before and after excluding F_1_ and F_2_ hybrids, in order to evaluate the potential contribution of hybridization to overall genetic variation. Specifically, we estimated the number of alleles (A), allelic richness (Ar), observed heterozygosity (H_O_), expected heterozygosity (H_E_), and nucleotide diversity (π) using the 5,702 SNP dataset. Genetic diversity was measured at both the population and regional levels. Number of alleles and allelic richness were estimated using the HIERFSTAT package v0.04-22 (Goudet, 2005) as implemented in R. Allelic richness was rarefied to a minimum of eight alleles (or four diploid samples). Meanwhile observed and expected heterozygosity were calculated using PLINK v1.90b6.12 (Purcell *et al*., 2007), and nucleotide diversity and the percentage of SNPs with missing data (NA) with VCFtools. Kruskal-Wallis tests and *post hoc* pairwise Wilcoxon tests were used to compare the levels of each diversity estimate among zones (allopatry, NC, NW).

#### Inter- and intraspecific genetic differentiation across hybrid zones

We assessed whether hybridization has led to changes in inter- or intraspecific genetic differentiation in sympatry. We compared overall interspecific genetic differentiation between *I. elegans* and *I. graellsii* across all three contexts—allopatry, the NC hybrid zone, and the NW hybrid zone—excluding F_1_ and F_2_ hybrids. To assess intraspecific genetic differentiation, we performed pairwise comparisons between localities across the same three contexts in both *I. elegans* and *I. graellsii* excluding F_1_ and F_2_ hybrids. All estimates were calculated using the 5,702 SNPs dataset by calculating F_ST_ (Weir and Cockerham, 1984) with 10,000 bootstraps with the R package StAMPP v1.6.1 (Pembleton *et al*., 2013). In the pairwise tests of population differentiation we employed the Bonferroni correction to control for multiple comparisons testing.

## RESULTS

### Reproductive isolation in the NC hybrid zone

#### Contribution of pre- and postmating barriers

In the NC hybrid zone, total prezygotic isolation was high in both directions, although slightly asymmetric (99.1% for *I. elegans* males × *I. graellsii* females; 80.1% for the reciprocal cross; Fig. 1A). Comparison with allopatry indicates that isolation in the *I. elegans* male × *I. graellsii* female cross has remained largely unchanged or only partially relaxed (100% in allopatry), unlike in the NW hybrid zone, where isolation in this direction is noticeably reduced relative to allopatry (45.9%). In contrast, the reciprocal cross (*I. graellsii* males × *I. elegans* females) shows stronger isolation relative to allopatry (8.2%). Similarly, in the NW hybrid zone the same cross direction also exhibited strong reproductive isolation (84.1%).

**Figure 1.**
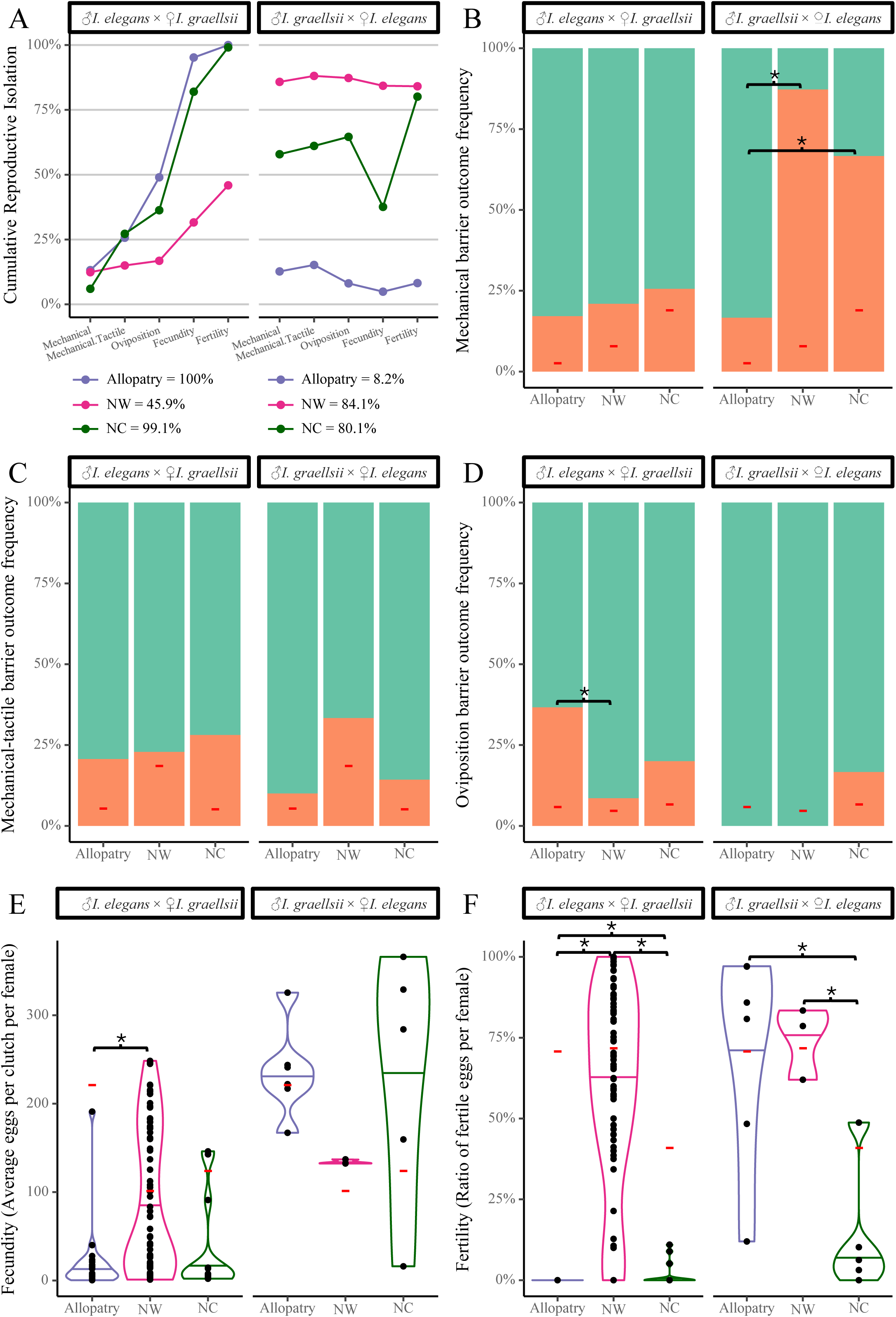
Prezygotic reproductive isolation between *Ischnura elegans* and *Ischnura graellsii*. Panel A shows cumulative strength of five prezygotic reproductive barriers in the NC hybrid zone (green), the NW hybrid zone (red) and allopatric populations (purple). Values at the bottom show the total cumulative isolation per zone for each type of cross. Panels B, C, and D show proportion of successful (top, green) and unsuccessful (bottom, orange) reproductive events for the mechanical (B), mechanical– tactile (C), and oviposition (D) barriers. Panels E and F, show violin plots of the distribution of fecundity (E) and (fertility F) per mated female. Horizontal red lines represent the mean values for conspecific crosses, used as a baseline of reproductive success (see Table 1). Asterisks (*) indicate statistically significant post hoc differences between zones (p < 0.05).

In NC hybrid zone, premating barriers in the *I. graellsii* male × *I. elegans* female cross were stronger (61.1%; Fig. 1A) than the premating isolation from this cross in allopatry (15.2%). Meanwhile, postmating barriers contributed strongly (71.9%) to isolation in the other cross direction (*I. elegans* male × *I. graellsii* female). In NW hybrid zone, premating barriers were low in the *I. elegans* male × *I. graellsii* female cross (15.0%) but very high in the reciprocal (88.1%; Fig. 1A), while postmating barriers were weaker (30.9% and -4%, respectively).

#### Reproductive success across reciprocal crosses and zones

The full GLM (reproductive isolation ∼ distribution + crosses + distribution*crosses) was selected as the most parsimonious model for the mechanical isolation, fecundity, and fertility barriers, whereas the GLM without the interaction term was the most parsimonious for the oviposition barrier. The mechanical-tactile isolation was independent of crosses and distribution (null model; Table 1).

In crosses between *I. graellsii* males and *I. elegans* females, the mechanical isolation was significantly stronger in both hybrid zones compared to allopatry, with the effect most pronounced in NW hybrid zone (Tukey test; NW vs. allopatry: p = 0.0002; NC vs. allopatry: p = 0.0289; Fig. 1B; Tables 1 & S2). In this cross direction we also detected significantly lower fertility (stronger isolation) in the NC hybrid zone than in both NW and allopatry (p < 0.0001; Fig. 1F; Tables 1 & S2).

In the reciprocal cross (*I. elegans* males × *I. graellsii* females), the only significant difference between hybrid zones was in fertility, which was lower (stronger isolation) in the NC hybrid zone (p < 0.0001; Figs. 1B–F; Tables 1 & S2). Additionally, we detected significantly higher fertilities (weaker isolation) in the NC hybrid zone than the allopatric crosses (p < 0.0001; Fig. 1F; Tables 1 & S2). The NW hybrid zone showed low isolation due to postmating barriers, with significantly weaker isolation due to oviposition (p = 0.0025), fecundity (p = 0.0178), and fertility (p < 0.0001; Figs. 1D–F; Tables 1 & S2) than the allopatric crosses.

### Genetic consequences of hybridization across hybrid zones

#### Complete set of SNPs and diagnostic SNPs

A total of 5,702 SNPs were detected after filtering. Of these, 2,127 (37.3%) were polymorphic in *I. elegans* but fixed in *I. graellsii*, and 1,711 (30.0%) were polymorphic in *I. graellsii* but fixed in *I. elegans*. Of the remaining SNPs, 1,421 (24.9%) were polymorphic in both species, 62 (1.1%) were fixed for the same allele in allopatric populations of both *I. elegans* and *I. graellsii,* and 381 (6.7%) species-specific (i.e., alternatively fixed between *I. elegans* and *I. graellsii* individuals from the allopatric populations; Table S3).

#### Genetic structure analyses

Genetic structure analyses based on the full dataset of 5,702 SNPs revealed consistent patterns of differentiation between *I. graellsii* and *I. elegans*, as well as varying degrees of admixture across the two hybrid zones.

Unsupervised ADMIXTURE analyses identified *K* = 2 as the most likely number of genetic clusters, based on the lowest cross-validation (CV) error, although CV values for *K* = 2 and *K* = 3 were similar (Fig. S3). At *K* = 2, clusters corresponded closely to *I. graellsii* and *I. elegans* (Fig. 2A). At *K* = 3, a third cluster emerged, separating *I. elegans* individuals from the NW hybrid zone from those in allopatry and the NC hybrid zone (Fig. 2A). For both *K* = 2 and *K* = 3, several individuals from the hybrid zones showed clear signs of admixture, with shared ancestry between clusters.

**Figure 2.**
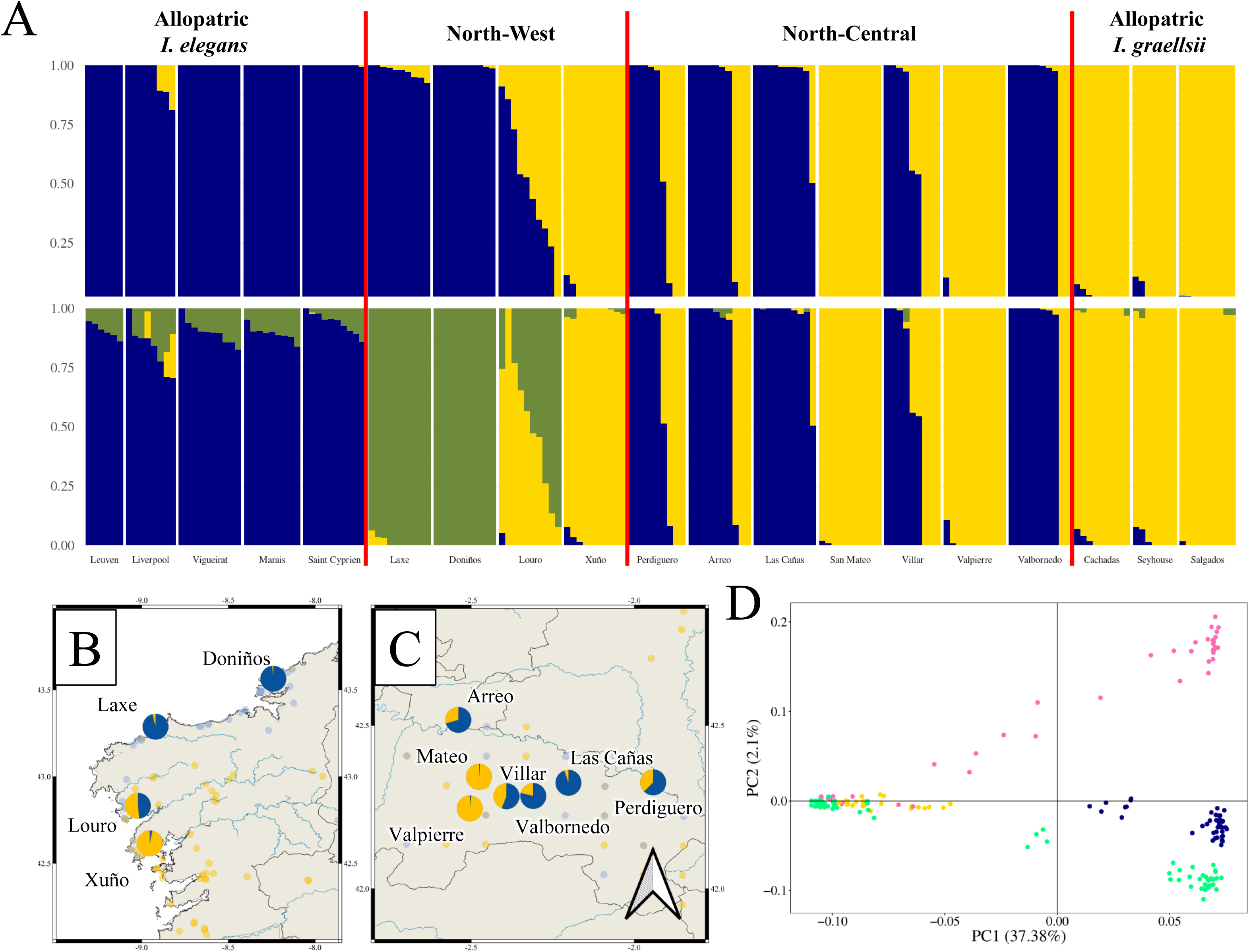
Genetic structure and diversity in *I. elegans*, *I. graellsii*, and their hybrids. Panel A presents ADMIXTURE results for K = 2 and K = 3, based on 5,702 SNPs without supervision. Panels B and C show the relative frequency of the species-specific alleles of the diagnostic SNPs for *I. elegans* (blue) and *I. graellsii* (yellow) across localities in the B) NW and C) NC hybrid zones. Panel D shows the Principal Component Analysis (PCA) using all SNPs. PC1 explains ∼37% and PC2 explains ∼2% of the total variance. Color coding indicates the *I. graellsii* allopatric (yellow), *I. elegans* allopatric (blue), NW hybrid zone (pink), and NC hybrid zone (green) regions.

PCA of the full SNP dataset provided further support for distinct species clusters and hybrid structure (Fig. 2D). The first principal component (PC1), which explained ∼37% of the total genetic variation, separated individuals of *I. elegans* and *I. graellsii* from allopatric populations, while individuals from hybrid zones varied in their positions along PC1, with many clustering closely with one of the parental species, while others occupied intermediate positions, consistent with admixed ancestry. The second principal component (PC2), explaining ∼2% of the variation, distinguished among *I. elegans* individuals from allopatry, the NC hybrid zone, and the NW hybrid zone. These show the presence of genetically pure individuals in both hybrid zones and a diversity of admixed genotypes. Introgression was also detected in Menorca in the Balearic Islands (Fig. S4.)

#### Admixture and assignment to hybrid classes

Using 381 diagnostic SNPs, we classified individuals from the NW and NC hybrid zones into eight hybrid classes based on hybrid index (HI) and heterozygosity (HET).

Individuals identified morphologically as *I. elegans* corresponded genetically either to pure *I. elegans* or to introgressed *I. elegans*. In contrast, individuals classified in the field as *I. graellsii* encompassed a much broader genetic range, including not only pure and introgressed *I. graellsii*, but also F_1_ hybrids, backcrosses with *I. graellsii*, and backcrosses with *I. elegans* (Table S4).

In both hybrid zones, ongoing hybridization and introgression were detected, but with markedly different patterns. In the NC hybrid zone, a large majority of individuals (89.7%, n=68) were genetically pure (*I. elegans* or *I. graellsii*) with a low proportion of admixed individuals. These hybrids were mostly F_1-_F_2_ hybrids, *I. graellsii* backcrosses, and introgressed *I. graellsii*, indicating limited and unidirectional introgression toward *I. graellsii* (Fig. 3A; Table S4). In contrast, the NW hybrid zone exhibited a more complex admixture structure. Although pure individuals (*I. elegans* or *I. graellsii*) were still the majority (65%, n=40), admixed genotypes were more frequent and diverse, including F_1_-F_2_ hybrids, backcrosses to both parental species, and introgressed of both *I. graellsii* and *I. elegans* (Fig. 3B; Table S4). This pattern suggests more extensive and bidirectional introgression in the NW hybrid zone. Consistent with this, we found statistical differences in hybrid class frequencies between zones (χ²(6) = 15.824, *p* = 0.0147), driven by a higher proportion of introgressed *I. elegans* individuals in the NW hybrid zone compared to the NC hybrid zone (Z = 4.536, *p* = 0.0331). Differences in hybridization patterns were also evident at the population scale with contrasts in both the direction and intensity of introgression, as well as in the relative frequency of the species-specific alleles of the diagnostic SNPs (Figs. 2B & C). In the NC hybrid zone, we found unidirectional introgression toward *I. graellsii* in two populations (Perdiguero, Valpierre) and ongoing hybridization (F_1_ and F_2_ hybrids, and backcrosses) in four populations (Cañas, Perdiguero, Villar, and Arreo; Fig. S4). In contrast, in the NW hybrid zone we observed bidirectional introgression in three populations (Doniños, Laxe, and Xuño) and ongoing hybridization (F_1_ and F_2_ hybrids, and backcrosses) in one population (Louro; Fig. S4). However, it is important to note that the sample sizes per locality were relatively small (n=10), which may influence the robustness of these findings.

**Figure 3.**
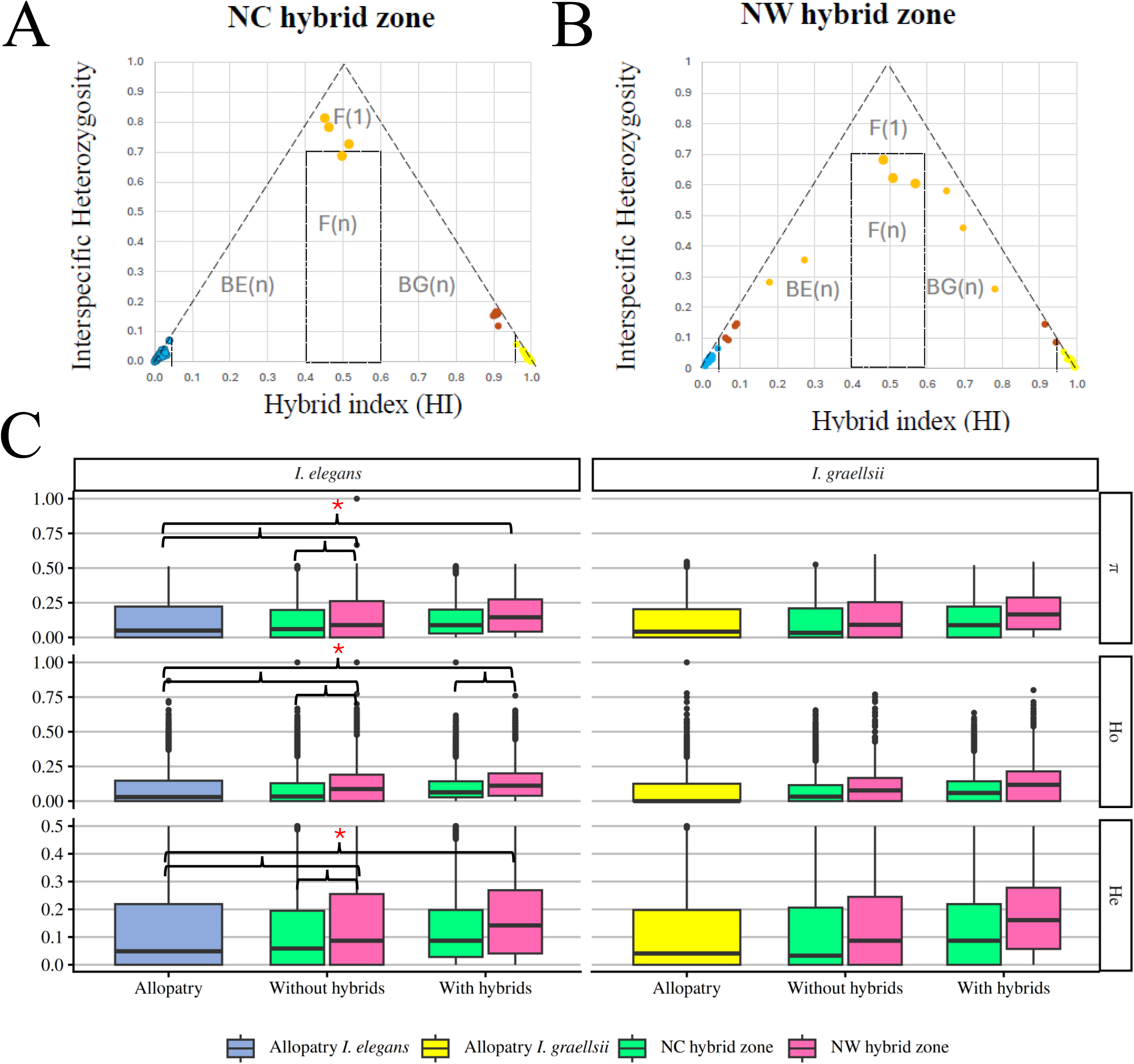
Ancestry proportions and hybrid classifications of *I. elegans* and *I. graellsii*. Panels A and B show individual estimates of hybrid index (HI) and interspecific heterozygosity (HET) calculated with INTROGRESS, based on 381 fixed SNPs, for individuals sampled in the A) NC hybrid zone and B) NW hybrid zone. F1 and F2 hybrids (orange) are located near the apex of the triangle, with high heterozygosity and intermediate ancestry values. Backcrosses to either *I. elegans* or *I. graellsii* (also orange) occupy positions with intermediate heterozygosity and asymmetric ancestry. Individuals with signs of introgression but low heterozygosity (brown) are positioned between pure individuals and backcrosses. Pure *I. elegans* (blue) and pure *I. graellsii* (yellow) cluster at opposite ends of the hybrid index axis. Panel C shows box plots of genetic diversity metrics across groups: nucleotide diversity (π), observed heterozygosity (Ho), and expected heterozygosity (He) with and without F1/F2 hybrids. Colors represent allopatric *I. elegans* (blue), allopatric *I. graellsii* (yellow), NC hybrid zone (green), and NW hybrid zone (pink) regions. Asterisks (*) indicate statistically significant differences between zones (p < 0.05).

#### Genetic diversity across hybrid zones

We examined patterns of genetic diversity to assess how hybridization and introgression influence variation in each hybrid zone. In *I. elegans*, excluding F_1_ and F_2_ hybrids, allelic richness, observed heterozygosity, expected heterozygosity, and nucleotide diversity were all significantly higher in the NW hybrid zone than in the NC hybrid zone or in allopatric populations (Fig. 3C; Tables S5–S7). When including F_1_ and F_2_ hybrids, only observed heterozygosity remained significantly higher in NW compared to NC hybrid zone, although all diversity measures were still higher in NW than in allopatry. In contrast, *I. graellsii* showed stable diversity across zones. The only significant difference observed was higher allelic richness in the NC hybrid zone compared to allopatry when hybrids were included (Fig. 3C; Tables S5–S7).

#### Inter- and intraspecific genetic differentiation across hybrid zones

Patterns of interspecific genetic differentiation, estimated using F_ST_, indicate that differentiation between *I. elegans* and *I. graellsii* tends to be lower in the NW hybrid zone (F_ST_ = 0.625) than in the NC hybrid zone (F_ST_ = 0.731), suggesting a trend toward more extensive introgression in the NW hybrid zone (Fig. 3). While formal statistical tests for these overall F_ST_ values are not available, pairwise intraspecific F_ST_ comparisons are generally consistent with this pattern: NW hybrid zone *I. elegans* populations show somewhat stronger differentiation (F_ST_ = 0.012–0.100; all 3/3 pairwise comparisons significant; Table S8) than NC hybrid zone populations (F_ST_ = 0.002–0.013; 3/10 pairwise comparisons significant; Table S8). In contrast, *I. graellsii* populations exhibit relatively low and homogeneous differentiation across zones, with pairwise F_ST_ ranging from 0–0.068 and fewer significant comparisons (Tables S9).

## DISCUSSION

Whether outcomes of hybridization are consistent across hybrid zones of the same species—and how they relate to the strength of reproductive isolation—are central questions in evolutionary biology. Reproductive isolation levels and the genetic consequences of hybridization differed markedly between the two hybrid zones that we studied.

We observed asymmetric reproductive productive barriers between reciprocal crosses in the NC hybrid zone. This was consistent with our prediction: we detected a stronger mechanical barrier in the NC hybrid zone than in allopatry in crosses between *I. graellsii* males and *I. elegans* females, mirroring the reinforcement process previously described for the NW hybrid zone (Arce-Valdés *et al*., 2024). However, we detected higher levels of postmating isolation (lower egg fertilities) in both reciprocal crosses in the NC hybrid zone than the NW hybrid zone (Fig. 1). These results indicate that pre- and postmating barriers evolve in a direction-specific manner in both hybrid zones, with NC hybrid zone retaining stronger ancestral reproductive isolation while NW hybrid zone exhibits more relaxed postmating barriers. Negative selection against maladaptive hybrids is expected to purge the Bateson-Dobzhansky-Müller (BDM) genetic incompatibilities responsible for postmating and postzygotic isolation (Turelli *et al*., 2014). However, the more recent formation of the NC hybrid zone than the NW hybrid zone suggests that BDM incompatibilities responsible for gametic incompatibility might still be present. Even more, the possible inflow from *I. elegans* allopatric individuals could be supplementing the NC hybrid zone with incompatibilities circumventing their purging. This would be evidenced by the absence of genetic structure between the *I. elegans* organisms from the NC hybrid zone and the allopatric populations of *I. elegans* (Fig. 2A). This contrasts with the incompatibilities purging suggested to have happened in the NW hybrid zone (Arce-Valdés *et al*., 2024), and the independent genetic cluster that *I. elegans* has evolved in the NW hybrid zone (Fig. 2A).

Genetically, in the NW hybrid zone we observed extensive admixture and bidirectional introgression. This caused increased intraspecific differentiation and reduced interspecific differentiation in comparison to the NC hybrid zone; and higher genetic diversity than in allopatry and the NC hybrid zone. In contrast, the NC hybrid zone showed stronger species boundaries detected as more limited and largely unidirectional gene flow, as well as only marginal differences in intra- or interspecific genetic differentiation, and genetic diversity with allopatry. Interestingly, while prezygotic barriers acted more asymmetrically in the NW hybrid zone than in the NC hybrid zone (Fig. 1A); introgressed individuals were detected towards both *I. elegans* and *I. graellsii* in the NW hybrid zone, but only towards *I. graellsii* in the NC hybrid zone (Figs. 3A & B). The introgression patterns of the NW hybrid zone are consistent with the levels of postzygotic isolation of the region, where hybrids formed in crosses between *I. elegans* males and *I. graellsii* females have been observed to successfully reproduce either with *I. elegans* or *I. graellsii* (Arce-Valdés *et al*., 2024). While future research could assess the reproductive fitness of hybrids from the NC hybrid zone to measure the postzygotic barriers in that area, the observed unidirectional introgression is consistent with the unidirectionally-inherited BDM incompatibilities (Turelli and Moyle, 2007) predicted to be present in X chromosome of *I. elegans* (Swaegers, Sánchez-Guillén, Chauhan, *et al*., 2022; Arce-Valdés *et al*., 2024). These regional differences concur with patterns found in other comparative hybrid zone studies that show that variation in hybridization outcomes often reflects differences in the time since sympatry—which strongly shapes the opportunity for reinforcement and introgression to accumulate (e.g. Kronforst *et al*., 2007; Lemmon and Juenger, 2017; Liao *et al*., 2019)—and the strength and direction of reproductive barriers (Vines *et al*., 2003; Lepais *et al*., 2009; Mandeville *et al*., 2017).

### Evolutionary consequences of hybridization across contrasting hybrid zones

The genomic patterns observed across hybrid zones provide key insights into how historical factors shape hybridization dynamics. Although the NW hybrid zone may be older than the NC hybrid zone, this inference remains tentative because it is based on differences in historical field records rather than formal demographic estimates and could instead reflect uneven sampling effort. Confirming whether these contrasts truly reflect differences in demographic history will therefore require dedicated demographic analyses (e.g. Collin *et al*., 2021; Excoffier *et al*., 2021).

Even with this uncertainty, the genomic contrasts between zones are clear. The NW hybrid zone exhibits extensive admixture, bidirectional introgression, and reduced interspecific differentiation—patterns broadly consistent with a longer or more sustained history of hybridization. Comparable increases in admixture and genomic diversity with prolonged hybridization have been documented in other systems (Buerkle and Rieseberg, 2001; Good *et al*., 2008; Aboim *et al*., 2010; Hohenlohe *et al*., 2011; Gompert *et al*., 2014), supporting the idea that extended secondary contact progressively erodes species boundaries and promotes a more unimodal hybrid zone structure —that is, a distribution dominated by genetically intermediate or admixed individuals rather than two distinct parental clusters (Barton and Hewitt, 1985). These genomic signatures also align with classical hybrid zone theory, which predicts that initial admixture between divergent genomes temporarily elevates genetic diversity before drift and selection remove maladaptive combinations (Barton and Hewitt, 1985; Harrison, 1990; Buerkle and Rieseberg, 2001). Consistent with this framework, diversity in the NW hybrid zone is moderately—but not extremely—high relative to allopatry, as expected under a scenario of repeated hybridization followed by the homogenizing effects of drift and selection. In contrast, the younger NC hybrid zone retains the genetic characteristics of more recent or limited secondary contact: admixture is sparse, differentiation remains high, and genetic diversity has not increased, reflecting a reduced window for recombination and introgression.

A crucial step in interpreting these patterns is distinguishing hybridization from alternative explanations such as incomplete lineage sorting. Although incomplete lineage sorting can mimic introgression under some scenarios (Holder *et al*., 2001), the spatial clustering of admixture—rather than the geographically random distribution expected under incomplete lineage sorting (Wang *et al*., 2019)—strongly supports hybridization. Moreover, the presence of introgressed alleles towards *I. graellsii* in both regions, together with additional introgression towards *I. elegans* specifically in the NW hybrid zone, is incompatible with a pure incomplete lineage sorting scenario and instead reflects true gene flow. This introgression in the NW hybrid zone may also help explain the localized relaxation of postcopulatory barriers observed there by Arce-Valdés et al. (2024), consistent with repeated hybridization, backcrossing, and reduction of interspecific differentiation.

Beyond these genomic patterns, the contrasting outcomes between hybrid zones are further shaped by differences in the strength and direction of reproductive barriers. Across both hybrid zones, reproductive isolation is strongly asymmetric, with reinforced mechanical isolation specifically targeting the *I. graellsii* male × *I. elegans* female cross. This consistent directionality suggests that selection against low-fitness hybrids acts disproportionately on this mating cross, in line with theoretical expectations that reinforcement evolves asymmetrically when hybridization costs differ between reciprocal crosses or sexes (Servedio and Noor, 2003; Yukilevich, 2012). By contrast, the reciprocal cross (*I. elegans* males × *I. graellsii* females)—which is mechanically viable—shows substantial spatial variation: in the NC hybrid zone, prezygotic isolation remains as strong as in allopatry, whereas in the NW hybrid zone these barriers show clear signs of relaxation. This divergence in barrier strength mirrors the genomic differences between hybrid zones: where barriers remain high (NC), gene flow is minimal and species boundaries persist; where barriers erode over time (NW), introgression accumulates and differentiation decreases.

### Phenotypic dominance and asymmetric hybrid formation across hybrid zones

The hybrids detected in our dataset included F_1_ and F_2_ individuals, as well as backcrosses with both parental species. Individuals identified morphologically as *I. elegans* corresponded genetically either to pure *I. elegans* or to introgressed individuals with *I. elegans*. In contrast, those classified in the field as *I. graellsii* encompassed a much broader range of genomic compositions, including pure *I. graellsii*, introgressed *I. graellsii*, F_1_ hybrids, and backcrosses with both parental species. This asymmetry suggests that hybrids and admixed individuals predominantly exhibit a *graellsii-like* phenotype, indicating a consistent morphological bias toward *I. graellsii* regardless of their underlying genomic background. Such dominance patterns in hybrid phenotypes align with theoretical expectations of non-additive inheritance and asymmetric expression (Turelli and Orr, 2000; Mallet, 2005) and are consistent with empirical evidence showing that dominance and parent-biased trait expression are pervasive in hybrids (Thompson *et al*., 2021; Runemark *et al*., 2025). This pattern is likely stronger in the NW hybrid zone by the unidirectional nature of hybridization, in which hybrid crosses occur primarily between *I. elegans* males and *I. graellsii* females. In later generations, mating continues to be biased: *I. elegans* males and hybrid females, as well as hybrid males with both *I. graellsii* and hybrid females (see Arce-Valdés *et al*., 2024). Importantly, *I. elegans* females were not involved in any hybrid crosses, because mechanical barriers prevent successful copulation with *I. graellsii* or hybrid males. Consistently, laboratory crosses reveal that F_1_ hybrids resemble *I. graellsii* in morphology, regardless of the maternal species (Sánchez-Guillén *et al*., 2005). This finding suggests that the *graellsii-like* phenotype is not simply the result of maternal effects but rather reflects a more intrinsic developmental or genetic dominance of *I. graellsii* traits in hybrid individuals.

### Conclusions

Comparing these two hybrid zones illustrates that reproductive isolation evolves in a highly context-dependent manner and that hybridization outcomes cannot be fully predicted from the genetic architecture of the species alone. Although our study focuses on *Ischnura* damselflies, the principles are likely general: (1) asymmetry and spatial variation in barrier strength strongly influence hybridization dynamics, (2) the direction and extent of introgression depend on demographic context and hybrid fitness, and (3) phenotypic dominance can obscure the underlying genomic composition of admixed individuals. These findings underscore the importance of considering both genomic composition and phenotypic expression when assessing hybridization outcomes, and they contribute to broader discussions about dominance, asymmetry, and predictability in hybrid zones. Understanding these dynamics is increasingly relevant in the context of climate change and anthropogenic range shifts, which create new zones of secondary contact and alter hybridization outcomes. Finally, our results highlight the value of integrating genomic, phenotypic, and ecological data to fully characterize hybrid zones, introgression patterns, and the evolution of reinforcement across taxa.

## ACKNOWLEDGEMENTS

We would like to thank the reviewers for their careful review and helpful comments, which have contributed to improving the quality of our manuscript. We are very grateful to Adolfo Cordero Rivera, who kindly allowed us to use his laboratory and material for the rearing experiments, and to the Zalandrana Odonatology group who kindly helped us with sampling and permitting in North-central Spain. We thank the following colleagues for kindly helping with collecting/sending samples: Adolfo Cordero Rivera, Iñaki Mezquita, Mario García-París, Bernat Garriós, Pere Luque, Xoaquín Baixeras, Francisco Cano, Jean Pierre Boudot, Jürgen Ott, Cedrick Vanappelghem, Philippe Lambret, and Phill Watts. We are grateful to Janet Nolasco Soto and Emmanuel Villafán de la Torre for technical support. Bioinformatics analyses were performed with the Huitzilin 2.0 HPC system at the Instituto de Ecología A.C. (INECOL). The research was funded by the European Union’s Marie Skłodowska-Curie Fellowship program (624538 and 753766 to JS, RAS-G, MW and BH), the Swedish Research Council (2016-00689 and 2022-04996 to BH) and the Mexican CONAHCYT (282922 to RAS-G).

## AUTHORS**’** CONTRIBUTIONS

RAS-G conceived the study idea. RAS-G, AB-G, LRA-V, JRC-R, JW acquired data. RAS-G, LRA-V and JS performed analyses. RAS-G wrote the first draft which was then revised and edited by all co-authors.

## DATA ARCHIVING

Raw sequencing data files were uploaded to the NCBI Sequence Read Archive: https://www.ncbi.nlm.nih.gov/bioproject/951651. The final filtered VCF input file as well as all the scripts for the full pipeline analysis were deposited on OSF at: https://osf.io/5kg87/?view_only=438667bce73d41ecab7137a65c625ded

