## Supplementary for "Hybridization outcomes reflect context-dependent reproductive isolation in two damselfly hybrid zones"

**Table of Contents:**

| **Figure S1.** Q-Q plots for the reproductive barriers GLMs. | Page 2 |
| --- | --- |
| **Figure S2.** Exploratory admixture plots. | Page 3 |
| **Figure S3.** CV-error plots for two admixture analyses | Page 4 |
| **Figure S4.** Comparison of several admixture plots. | Page 5 |
| **Table S1.** Sampled localities for molecular and reproductive barriers analyses. | Page 7 |
| **Table S2.** Absolute values of reproductive isolation per type of cross. | Page 8 |
| **Table S3.** Frequency of polymorphisms in allopatric samples | Page 9 |
| **Table S4.** Distribution of HI/HET values. | Page 10 |
| **Table S5.** Summary statistics of genetic diversity | Page 11 |
| **Table S6.** Statistical tests of genetic diversity between regions | Page 12 |
| **Table S7.** Genetic diversity pairwise statistical tests between regions | Page 13 |
| **Table S8.** Pairwise F_ST_ estimations between *I. elegans* populations | Page 14 |
| **Table S9.** Pairwise F_ST_ estimations between *I. graellsii* populations | Page 15 |
| **Supplemental Information References** | Page 16 |

| **Mechanical**  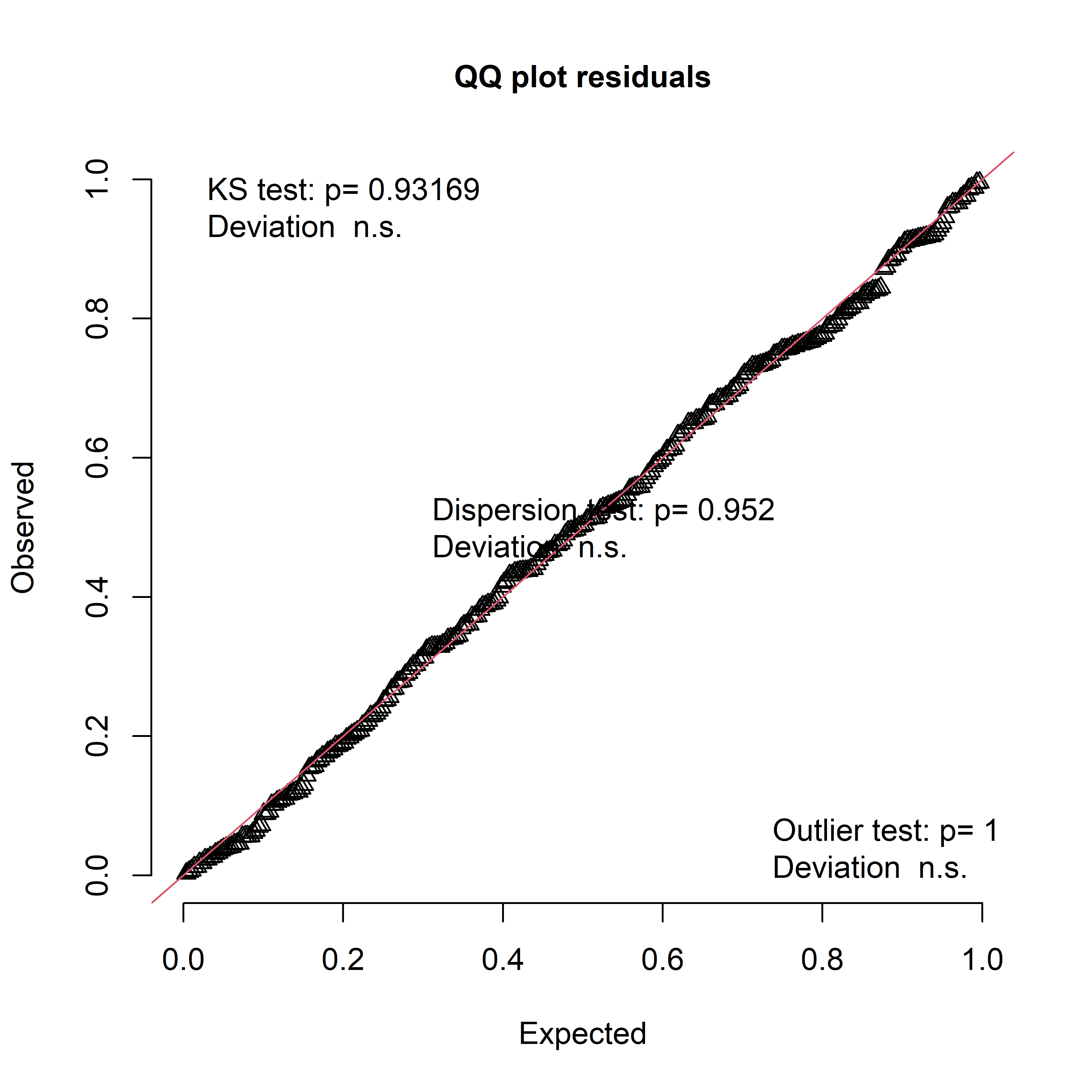 | **Mechanical-Tactile**  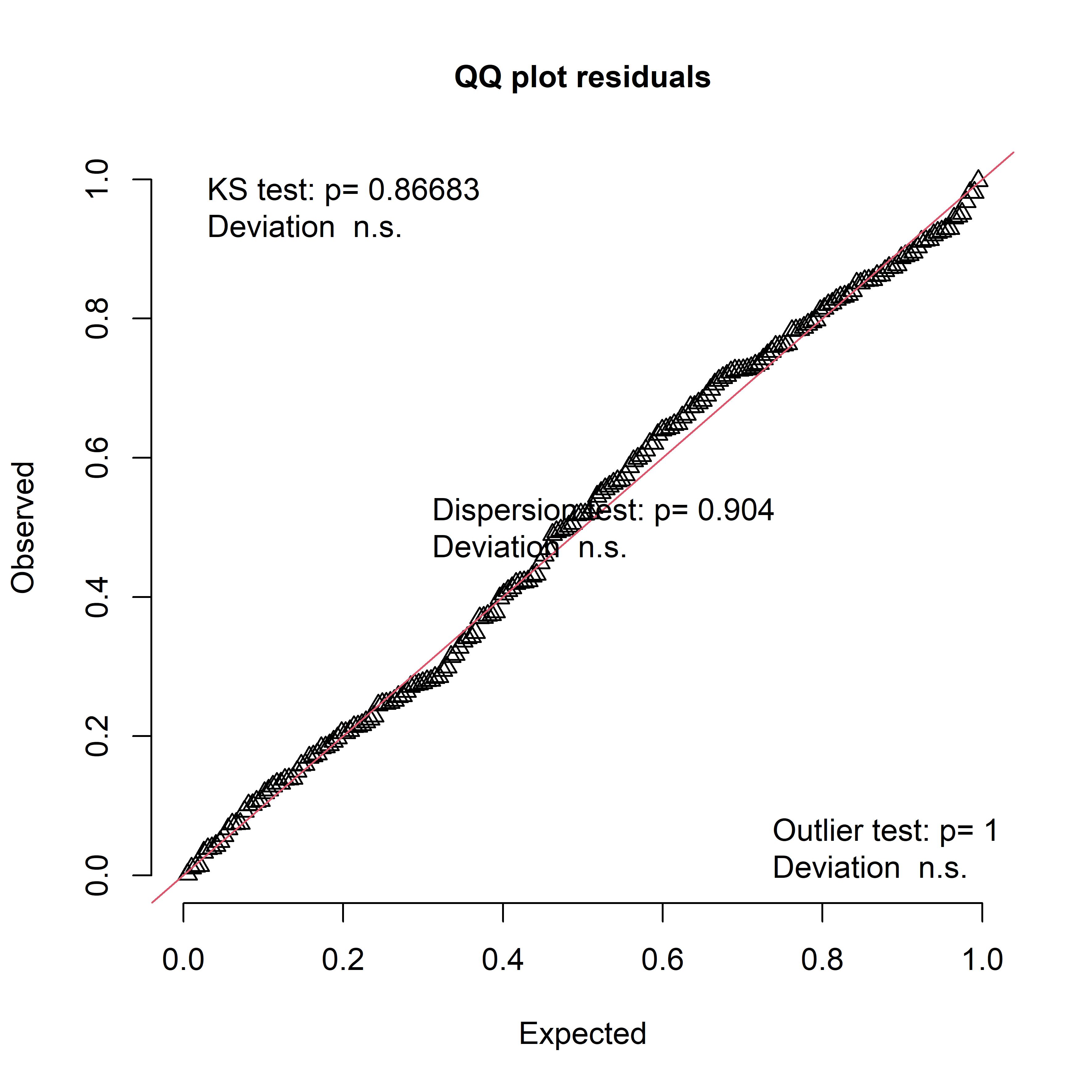 |
| --- | --- |
| **Oviposition**  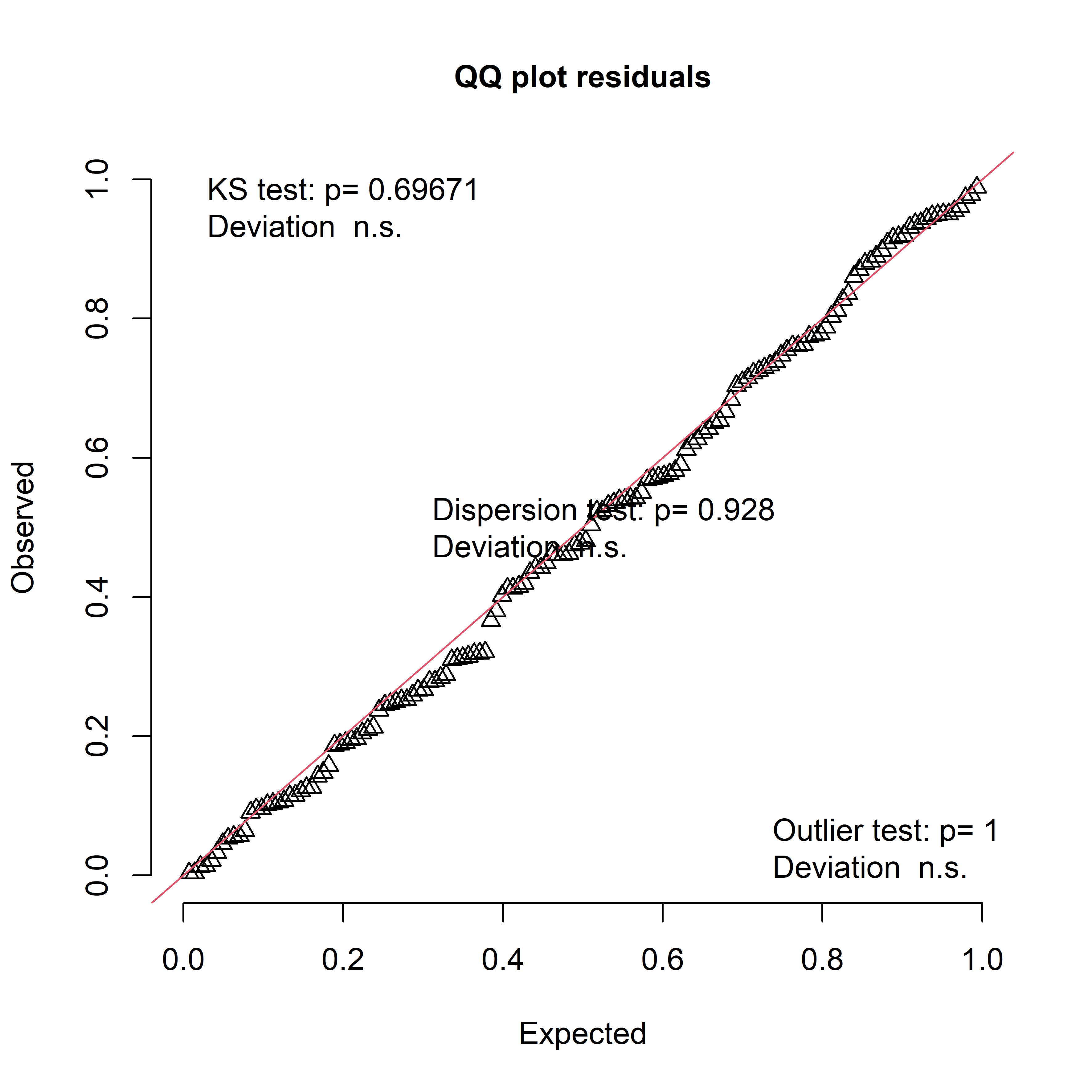 | **Fecundity**  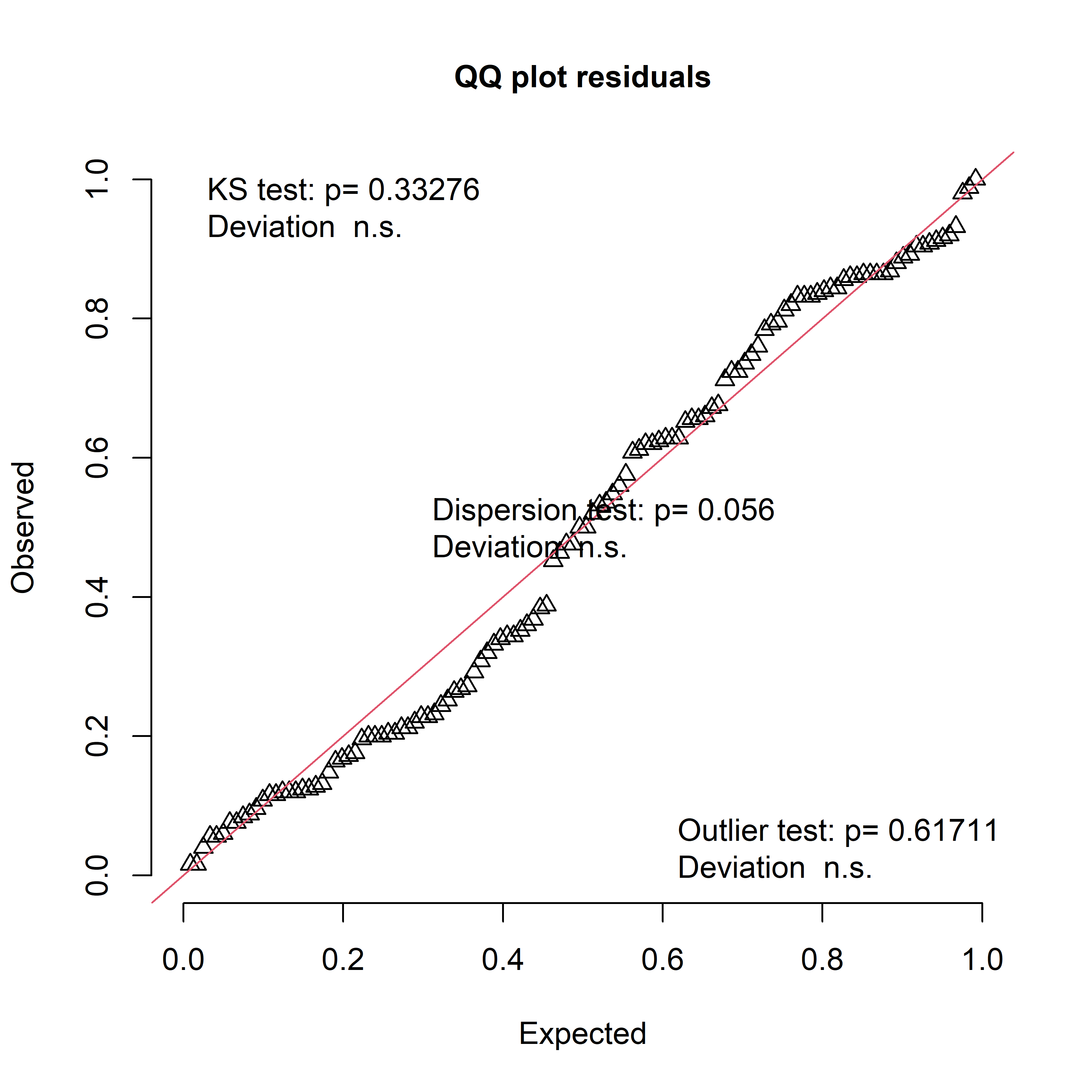 |
| **Fertility**  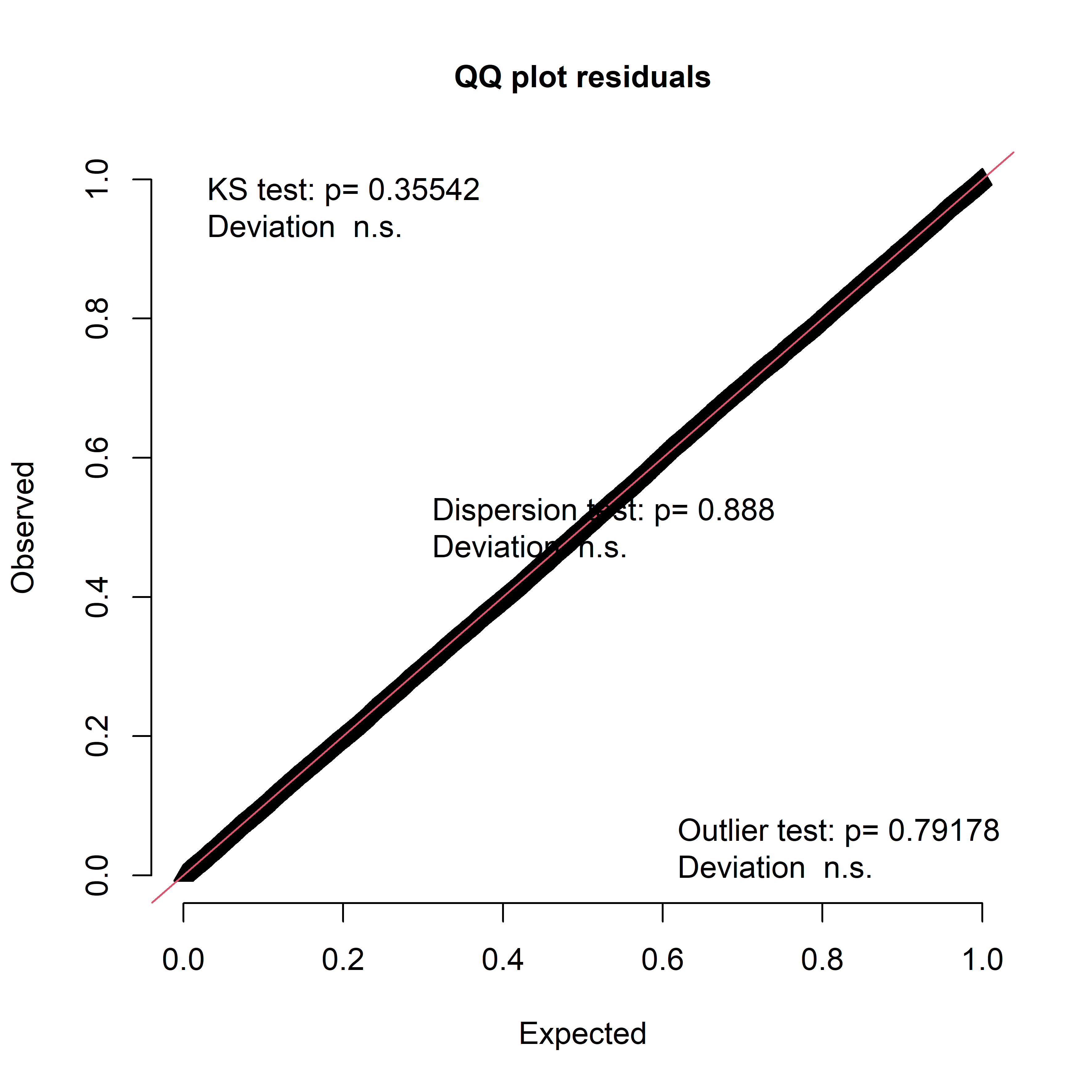 | |

**Figure S1.** Dharma Q-Qplots for the selected GLM of each of the measured reproductive barriers across allopatry and the NW and NC hybrid zones.

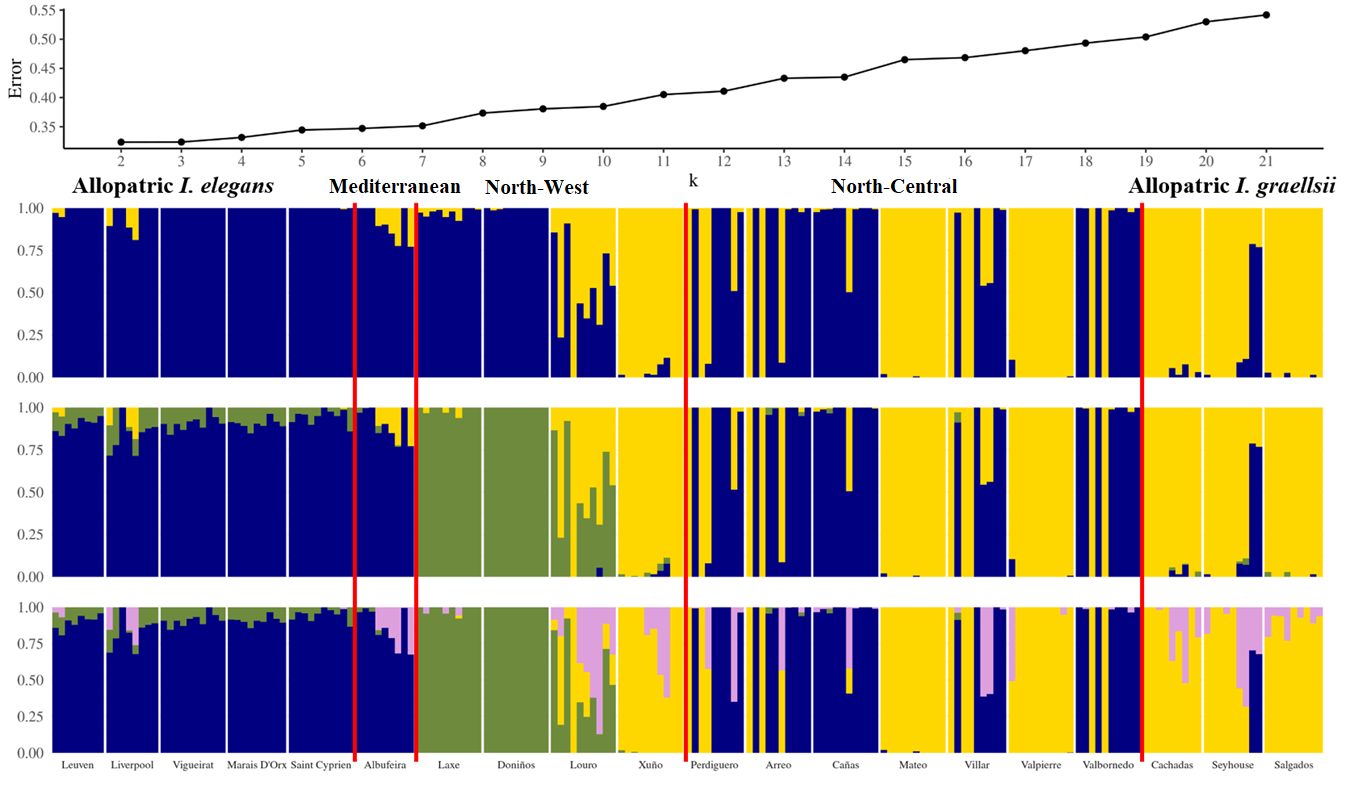

**Figure S2.** Unsupervised ADMIXTURE plots (*K* = 2, *K* = 3) and exploratory cross-validation error values for 189 *I. elegans*, *I. graellsii*, and hybrid individuals based on the full SNP dataset (5,702 SNPs). The analysis includes two replicate samples and two *I. graellsii* individuals from its allopatric range that show signals of possible hybridization with a third, unidentified *Ischnura* species (highlighted with a black box). These four samples were excluded from all subsequent analyses.

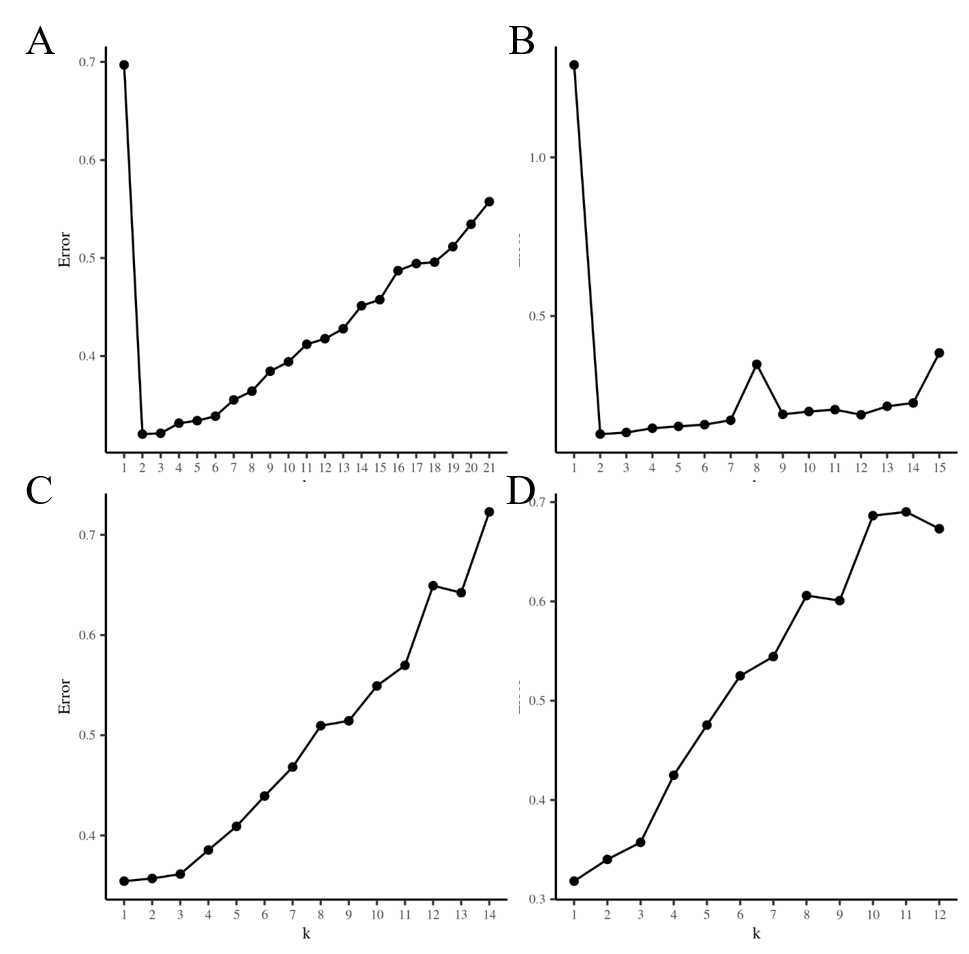

**Figure S3.** *CV-error* plots for different numbers of clusters (K) for **A)** All (5,702) SNPs from K = 1 to K = 21; **B)** Alternatively-fixed diagnostic SNPs (381) from K=1 to K=15 for all samples.

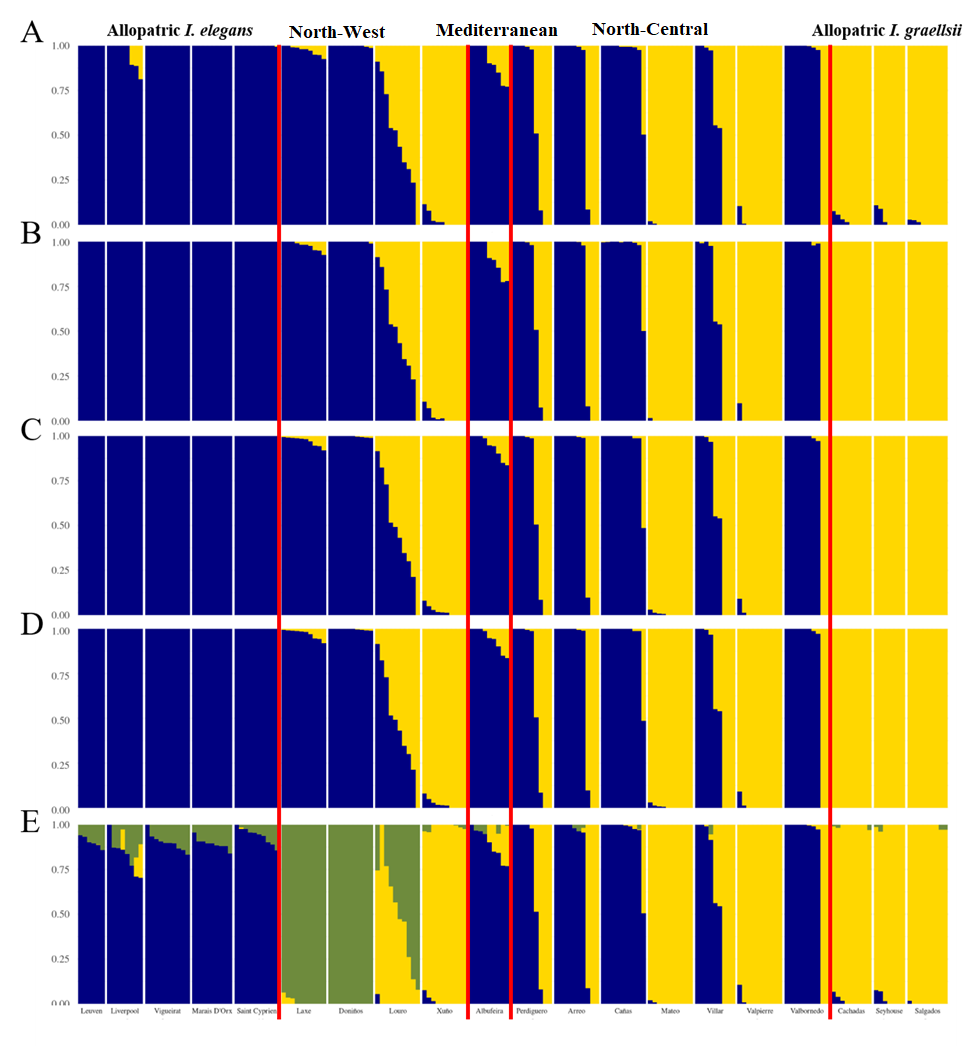

**Figure S4.** ADMIXTURE results for *K* = 2 and *K* = 3 using the full SNP dataset (5,702 SNPs) and the subset of alternatively fixed diagnostic SNPs (381 SNPs), under both unsupervised and supervised models. A) *K* = 2, full dataset, unsupervised; B)*K* = 2, full dataset, supervised; C) *K* = 2, diagnostic SNPs, unsupervised; D) *K* = 2, diagnostic SNPs, supervised; E) K = 3, full dataset, unsupervised.

This analysis includes samples from a third potential hybrid zone totaling nine individuals, which were excluded from all subsequent analyses. Individuals from this Mediterranean locality (Albufeira, Menorca, Balearic Islands) exhibited strong introgression of *I. graellsii* alleles, despite the absence of pure *I. graellsii* individuals or clear F₁/F₂ hybrids. This is unexpected given that historical records (1910–1991) report the exclusive presence of *I. elegans* in the Balearic Islands (Riservato et al., 2009), a finding supported by a recent doctoral thesis focused on Menorca (Soler Monzó, 2015). Interestingly, although individuals from Menorca have been phenotypically identified as *I. elegans*, the frequency of female morphs differs from other Mediterranean populations. In particular, the *infuscans-obsoleta* morph is predominant (Sánchez-Guillén et al., 2011), suggesting either stochastic processes during colonization or the assimilation of *I. graellsii*-like morph frequencies through hybridization.

**Table S1.** Hybrid zone, species proportions —estimated as the proportion of males of each species captured during previous sampling events, which ranged from one to five per population—, sampling year, country, latitude and longitude for each sampling locality. Sample size for molecular analysis (N) indicates the number of morphologically identified samples (*I. elegans*-like (I. e) and *I. graellsii*-like (I. g).

| **Zone** | **Species proportions** | | **Sampling locality, country** | **N** | **Year** | **Latitude** | **Longitude** |
| --- | --- | --- | --- | --- | --- | --- | --- |
|  | ***I. graellsii*** | ***I. elegans*** |  |  |  |  |  |
| Allopatry | 100% |  | ^†^Seyhouse, Algeria | 9 (I. g) | 2011 | 36.2827 | 7.2227 |
|  | 100% |  | Salgados, Portugal | 9 (I. g) | 2007 | 37.0971 | -8.3344 |
|  | 100% |  | Cachadas, Spain | 9 (I. g) | 2015 | 42.2712 | -8.5055 |
|  |  | 100% | Leuven, Belgium | 6 (I. e) | 2015 | 50.5341 | 4.4318 |
|  |  | 100% | Liverpool, UK | 8 (I. e) | 2008 | 53.2439 | -2.5840 |
|  |  | 100% | Vigueirat, France | 10 (I. e) | 2014 | 43.5311 | 4.3012 |
|  |  | 100% | Marais D’Orx, France | 9 (I. e) | 2015 | 43.5792 | -1.3982 |
|  |  | 100% | Saint Cyprien, France | 10 (I. e) | 2015 | 42.3823 | 3.0112 |
| North-Central  hybrid zone | 95-100% | 0-5% | Mateo, Spain | 10 (I. g) | 2015 | 42.3036 | -2.5154 |
|  | 95-100% | 0-5% | Valpierre, Spain | 10 (I. g) | 2015 | 42.2760 | -2.5436 |
|  | 0-26% | 74-100% | Villar, Spain | 6 (I. e), 4 (I. g) | 2015 | 42.2440 | -2.4211 |
|  | 0-19% | 81-100% | Arreo, Spain | 7 (I. e), 3 (I. g) | 2008 | 42.4775 | -2.5787 |
|  | 10-19% | 71-90% | Perdiguero, Spain | 5 (I. e), 4 (I. g) | 2015 | 42.2864 | -1.9818 |
|  | 40-42% | 58-60% | Las Cañas, Spain | 7 (I. e), 2 (I. g) | 2015 | 42.2853 | -2.2414 |
|  | 19-68% | 32-81% | Valbornedo, Spain | 8 (I. e), 2 (I. g) | 2015 | 42.2449 | -2.3490 |
| North-West  hybrid zone |  | 100% | Doniños, Spain | 10 (I. e) | 2014 | 43.4900 | -8.3150 |
|  | 0-2% | 98-100% | Laxe, Spain | 10 (I. e) | 2014 | 43.2128 | -8.9937 |
|  | 63-98% | 2-38% | Louro, Spain | 1 (I. e), 9 (I. g) | 2013 | 42.7581 | -9.0951 |
|  | 100% |  | Xuño, Spain | 10 (I. g) | 2014 | 42.5413 | -9.0264 |
| Mediterranean  (Menorca) |  | 100% | ^‡^Albufeira, Spain | 9 (I. e) | 2009 | 39.9495 | 4.2497 |
| ^†^Two samples removed after SNP calling due to possible hybridization with a third unknown species.  ^‡^ADMIXTURE analyses to test past hybridization in the zone are given in the Supplementary Figure S2. | | | | | | | |

**Table S2.** Absolute values of reproductive isolation per type of cross and reproductive barrier for allopatry and the NC and NW hybrid zones. Values between parenthesis show the sample size of each measurement.

| **Zone** | **Type** | **Male** | **Female** | **Mechanical** | **Mechanical-Tactile** | **Oviposition** | **Fecundity** | **Fertility** |
| --- | --- | --- | --- | --- | --- | --- | --- | --- |
| Allopatry | Conspecifics | *I. elegans* | *I. elegans* | 0.005 (20) | 0.034 (19) | 0.039 (35) | -0.226 (31) | 0.025 (31) |
|  | Conspecifics | *I. graellsii* | *I. graellsii* | -0.005 (25) | -0.034 (24) | -0.039 (24) | 0.226 (23) | -0.025 (23) |
|  | Heterospecifics | *I. elegans* | *I. graellsii* | 0.132 (70) | 0.144 (58) | 0.313 (30) | 0.907 (19) | 1.000 (19) |
|  | Heterospecifics | *I. graellsii* | *I. elegans* | 0.127 (12) | 0.029 (10) | -0.085 (6) | -0.034 (6) | 0.035 (6) |
| NW | Conspecifics | *I. elegans* | *I. elegans* | 0.108 (46) | 0.048 (37) | 0.07 (38) | -0.074 (33) | -0.038 (33) |
|  | Conspecifics | *I. graellsii* | *I. graellsii* | -0.108 (12) | -0.048 (12) | -0.07 (16) | 0.074 (16) | 0.038 (16) |
|  | Heterospecifics | *I. elegans* | *I. graellsii* | 0.124 (105) | 0.03 (83) | 0.021 (82) | 0.178 (75) | 0.209 (75) |
|  | Heterospecifics | *I. graellsii* | *I. elegans* | 0.858 (47) | 0.161 (6) | -0.07 (3) | -0.235 (3) | -0.013 (3) |
| NC | Conspecifics | *I. elegans* | *I. elegans* | 0.069 (19) | 0.077 (14) | 0.021 (19) | -0.35 (17) | -0.113 (17) |
|  | Conspecifics | *I. graellsii* | *I. graellsii* | -0.069 (13) | -0.077 (11) | -0.021 (15) | 0.35 (14) | 0.113 (14) |
|  | Heterospecifics | *I. elegans* | *I. graellsii* | 0.06 (43) | 0.226 (32) | 0.125 (15) | 0.718 (12) | 0.951 (12) |
|  | Heterospecifics | *I. graellsii* | *I. elegans* | 0.579 (21) | 0.077 (7) | 0.088 (6) | -0.761 (5) | 0.682 (5) |

**Table S3.** Frequency of polymorphisms in both *Ischnura* species allopatric distributions.

| **Polymorphism type Count** | **Region** | **Count** | **Frequency** |
| --- | --- | --- | --- |
| Alternately fixed SNPs (Diagnostic SNPs) | Allopatry | 381 | 6.7% |
| Fixed for the same allele | Allopatry | 62 | 1.1% |
| *I. elegans* and *I. graellsii* shared polymorphisms | Allopatry | 1421 | 24.9% |
| SNP polymorphic in *I. elegans* but one allele fixed in *I. graellsii* | Allopatry | 2127 | 37.3% |
| SNP polymorphic in *I. graellsii* but one allele fixed in *I. elegans* | Allopatry | 1711 | 30.0% |
| Total SNPs analyzed | - | 5702 | 100% |

**Table S4.** Distribution of individuals in eight ranges of HI/HET values from Introgress. North-central hybrid zone included 68 samples (33 *I. elegans*-like and 35 *I. graellsii*-like) and the north-west hybrid zone included 40 samples (21 *I. elegans*-like and 19 *I. graellsii*-like). All hybrids were *a priori,* morphologically identified as *I. graellsii,* which is consistent with the morphology of the F_1_ and F_2_ hybrids from the laboratory (Monetti et al., 2002; Sánchez-Guillén et al., 2005) which usually resemble *I. graellsii* individuals.

| **Classes** | **Hybrid Index**  **(HI)** | **Heterozygosity**  **(HET)** | **Allopatry** | **Spanish hybrid zone** | |
| --- | --- | --- | --- | --- | --- |
|  |  |  |  | **North-central** | **North-west** |
| Morphologically *I. graellsii* | | | | | |
| Pure *I. graellsii* | <0.05 | ≤0.08 | 25 (100%) | 28 (80.00%) | 9 (47.37%) |
| Introgressed *I. graellsii* | 0.05-0.10 | ˂0.16 | - | 2 (5.71%) | 2 (10.53%) |
| Back to *I. graellsii* | 0.11-0.39 | ˂0.60 | - | 1 (2.86%) | 3 (15.79%) |
| F_1_ hybrids | 0.40-0.60 | ≥0.600 | - | 4 (11.43%) | 3 (15.79%) |
| Back to *I. elegans* | 0.61-0.89 |  |  | 0 | 2 (10.53%) |
| Morphologically *I. elegans* | | | | | |
| Introgressed *I. elegans* | 0.90-0.95 | ˂0.16 | - | 0 | 4 (19.05%) |
| Pure *I. elegans* | ˃0.95 | ≤0.08 | 43 (100%) | 33 (100%) | 17 (80.95%) |
| All sampled individuals | | | | | |
| Pure *I. graellsii* | <0.05 | ≤0.08 | 25 (100%) | 28 (41.17%) | 9 (22.50%) |
| Introgressed *I. graellsii* | 0.05-0.10 | ˂0.16 |  | 2 (2.94%) | 2 (5.00%) |
| Back to *I. graellsii* | 0.11-0.39 | ˂0.60 |  | 1 (1.47%) | 3 (7.50%) |
| F1 hybrids | 0.40-0.60 | ≥0.600 |  | 4 (5.88%) | 3 (7.50%) |
| Back to *I. elegans* | 0.61-0.89 | ˂0.60 |  | 0 | 2 (5.00%) |
| Introgressed *I. elegans* | 0.90-0.95 | ˂0.16 |  | 0 | 4 (10.00%) |
| Pure *I. elegans* | ˃0.95 | ≤0.08 | 43 (100%) | 33 (48.52%) | 17 (42.5%) |

**Table S5.** Summary statistics of genetic diversity per regions. Genetic diversity was measured in terms of: **n** = Sample size; **NA** = Percentage of SNPs with missing data; **A** = Number of alleles; **Ar** = Allelic richness rarefacted for four samples; **H_O_** = Observed heterozygosity; **H_E_** = Expected heterozygosity; and **π** = nucleotide diversity.

| **Region** | **Species** | **n** | **NA** | **A** | **Ar** | **H_O_** | **H_E_** | **π** |
| --- | --- | --- | --- | --- | --- | --- | --- | --- |
| Allopatric | *I. elegans* | 43 | 16.8% | 9250 | 7639.5 | 0.088 | 0.121 | 0.122 |
| North-west hybrid zone | *I. elegans* | 23 | 6.5% | 9884 | 8096.6 | 0.122 | 0.150 | 0.154 |
|  | *I. elegans* + hybrids | 26 | 7.2% | 10472 | 8482.3 | 0.136 | 0.171 | 0.175 |
|  | *I. graellsii* | 14 | 8.8% | 9432 | 8085.6 | 0.112 | 0.141 | 0.147 |
|  | *I. graellsii* + hybrids | 17 | 9.4% | 9999 | 8539.5 | 0.137 | 0.172 | 0.178 |
| North-central hybrid zone | *I. elegans* | 33 | 9.1% | 9432 | 7600.4 | 0.085 | 0.117 | 0.119 |
|  | *I. elegans* + hybrids | 37 | 10.4% | 10430 | 7944.2 | 0.098 | 0.133 | 0.135 |
|  | *I. graellsii* | 31 | 8.1% | 8952 | 7447.7 | 0.079 | 0.110 | 0.112 |
|  | *I. graellsii* + hybrids | 35 | 9.6% | 9871 | 7865.6 | 0.093 | 0.131 | 0.133 |
| Allopatric | *I. graellsii* | 25 | 22.1% | 8774 | 7487.1 | 0.076 | 0.111 | 0.115 |

**Table S6. Statistical assessment of genetic diversity across geographic regions (two hybrid zones and allopatric populations).** Comparative analyses of genetic diversity metrics among regions were performed using the non-parametric Kruskal-Wallis rank sum test, independently applied to each diversity category. **A** = Number of alleles; **Ar** = Allelic richness rarefacted for four samples; **H_O_** = Observed heterozygosity; **H_E_** = Expected heterozygosity; and **π** = nucleotide diversity.

| **Estimate** | | ***I. elegans*** | ***I. elegans* + hybrids** | ***I. graellsii*** | ***I. graellsii + hybrids*** |
| --- | --- | --- | --- | --- | --- |
| A | χ^2^ | 4.141 | 6.251 | 4.844 | 7.262 |
|  | df | 2 | 2 | 2 | 2 |
|  | p | 0.126 | **0.044** | 0.089 | **0.026** |
| Ar | χ^2^ | 7.543 | 7.578 | 5.723 | 6.691 |
|  | df | 2 | 2 | 2 | 2 |
|  | p | **0.023** | **0.023** | 0.057 | **0.035** |
| H_O_ | χ^2^ | 6.488 | 7.543 | 6.575 | 5.108 |
|  | df | 2 | 2 | 2 | 2 |
|  | p | **0.039** | **0.023** | **0.0373** | 0.077 |
| H_E_ | χ^2^ | 7.543 | 6.312 | 4.844 | 3.330 |
|  | df | 2 | 2 | 2 | 2 |
|  | p | **0.023** | **0.043** | 0.089 | 0.189 |
| π | χ^2^ | 6.435 | 6.751 | 3.857 | 2.851 |
|  | df | 2 | 2 | 2 | 2 |
|  | p | **0.040** | **0.034** | 0.145 | 0.240 |
| **Bold** = Statistically significant (p < 0.05). | | | | | |

**Table S7. Pairwise statistical comparisons of genetic diversity among regions.**
Pairwise comparisons of genetic diversity between regions were conducted using the exact Wilcoxon rank sum test. Each zone was treated as an independent sample within its respective region to evaluate significant differences in diversity metrics. **A** = Number of alleles; **Ar** = Allelic richness rarefacted for four samples; **H_O_** = Observed heterozygosity; **H_E_** = Expected heterozygosity; and **π** = nucleotide diversity.

|  | | Allopatric Area | North-west hybrid zone | Allopatric Area | North-west hybrid zone |
| --- | --- | --- | --- | --- | --- |
|  | | ***I. elegans*** | | ***I. graellsii*** | |
| North-west hybrid zone | A | 0.143 | - | 0.200 | - |
|  | Ar | **0.036** | - | 0.200 | - |
|  | H_O_ | **0.036** | - | 0.200 | - |
|  | H_E_ | **0.036** | - | 0.200 | - |
|  | π | **0.036** | - | 0.200 | - |
| North-central hybrid zone | A | 0.841 | 0.071 | 0.667 | 0.056 |
|  | Ar | 0.222 | **0.036** | 0.267 | 0.056 |
|  | H_O_ | 0.841 | **0.036** | 0.067 | 0.111 |
|  | H_E_ | 0.222 | **0.036** | 0.667 | 0.056 |
|  | π | 1.000 | **0.036** | 0.830 | 0.110 |
|  | | ***I. elegans* + hybrids** | | ***I. graellsii* + hybrids** | |
| North-west hybrid zone | A | **0.036** | - | 0.200 | - |
|  | Ar | **0.036** | - | 0.200 | - |
|  | H_O_ | **0.036** | - | 0.200 | - |
|  | H_E_ | **0.036** | - | 0.200 | - |
|  | π | **0.036** | - | 0.200 | - |
| North-central hybrid zone | A | 0.151 | 0.250 | 0.067 | 0.056 |
|  | Ar | **0.032** | 0.393 | **0.033** | 0.222 |
|  | H_O_ | 0.222 | **0.036** | 0.067 | 0.500 |
|  | H_E_ | 0.222 | 0.143 | 0.380 | 0.330 |
|  | π | 0.095 | 0.250 | 0.380 | 0.500 |
| **Bold** = Statistically significant (p < 0.05). | | | | | |

**Table S8.** Pairwise F_ST_ estimations between *I. elegans* localities. F_ST_ values above the diagonal and p values below the diagonal. **Blue** = *I. elegans* allopatric region; **pink** = north-west hybrid zone; **green** = north-central hybrid zone; ***=**statistically significant p-values after Bonferroni correction (p < 0.05/78).

|  | Leuven | Liverpool | Vigueirat | Marais D’Orx | Saint Cyprien | Laxe | Doniños | Louro | Perdiguero | Arreo | Cañas | Villar | Valbornedo |
| --- | --- | --- | --- | --- | --- | --- | --- | --- | --- | --- | --- | --- | --- |
| Leuven | - | 0.0078 | **0.0193** | 0.0023 | **0.0211** | **0.0929** | **0.1023** | **0.1564** | **0.0838** | **0.0848** | **0.0913** | **0.0902** | **0.0915** |
| Liverpool | *0.0704* | - | **0.0447** | **0.0211** | **0.0378** | **0.0866** | **0.1040** | **0.1173** | **0.0718** | **0.0696** | **0.0726** | **0.0795** | **0.0815** |
| Vigueirat | *0** | *0** | - | **0.0142** | **0.0165** | **0.0961** | **0.0978** | **0.1696** | **0.0789** | **0.073** | **0.0772** | **0.0815** | **0.0799** |
| Marais D’Orx | *0.2526* | *0** | *0** | - | **0.0152** | **0.094** | **0.0966** | **0.1728** | **0.0839** | **0.0788** | **0.0791** | **0.0891** | **0.0851** |
| Saint Cyprien | *0** | *0** | *0** | *0** | - | **0.0951** | **0.0992** | **0.1687** | **0.059** | **0.0562** | **0.0592** | **0.0658** | **0.0616** |
| Laxe | *0** | *0** | *0** | *0** | *0** | - | **0.0123** | **0.0692** | **0.1138** | **0.1058** | **0.1106** | **0.1122** | **0.1152** |
| Doniños | *0** | *0** | *0** | *0** | *0** | *0** | - | **0.1005** | **0.1282** | **0.1164** | **0.1191** | **0.1289** | **0.1272** |
| Louro | *0** | *0** | *0** | *0** | *0** | *0** | *0** | - | **0.1585** | **0.1604** | **0.1684** | **0.1483** | **0.1846** |
| Perdiguero | *0** | *0** | *0** | *0** | *0** | *0** | *0** | *0** | - | 0.0021 | 0.0053 | 0.0111 | 0.0023 |
| Arreo | *0** | *0** | *0** | *0** | *0** | *0** | *0** | *0** | *0.3082* | - | 0.0043 | 0.0055 | **0.0117** |
| Cañas | *0** | *0** | *0** | *0** | *0** | *0** | *0** | *0** | *0.0947* | *0.0897* | - | 0.0132 | 0.0098 |
| Villar | *0** | *0** | *0** | *0** | *0** | *0** | *0** | *0** | *0.0278* | *0.1296* | *0.0036* | - | 0.0070 |
| Valbornedo | *0** | *0** | *0** | *0** | *0** | *0** | *0** | *0** | *0.304* | *0** | *0.0009* | *0.0539* | - |

**Table S9.** Pairwise F_ST_ estimations between *I. graellsii* localities. F_ST_ values above the diagonal and p values below the diagonal. **Yellow** = *I. graellsii* allopatric region; **pink** = north-west hybrid zone; **green** = north-central hybrid zone; ***=**statistically significant *p-values* after Bonferroni correction (p < 0.05/66).

|  | Cachadas | Seyhouse | Salgados | Louro | Xuño | Perdiguero | Arreo | Cañas | Mateo | Villar | Valpierre | Valbornedo |
| --- | --- | --- | --- | --- | --- | --- | --- | --- | --- | --- | --- | --- |
| Cachadas | - | **0.0678** | 0.0082 | **0.0947** | 0 | 0.0106 | **0.018** | 0 | **0.0189** | 0.0021 | **0.0097** | 0.0063 |
| Seyhouse | *0** | - | **0.0592** | **0.121** | **0.0722** | **0.058** | **0.0623** | 0.0098 | **0.0893** | **0.0835** | **0.0749** | **0.0569** |
| Salgados | *0.0456* | *0** | - | **0.1052** | **0.0146** | 0.0042 | 0.0177 | 0 | **0.0285** | 0 | **0.0178** | 0 |
| Louro | *0** | *0** | *0** | - | **0.0793** | **0.0544** | **0.0465** | 0 | **0.1404** | **0.0415** | **0.1182** | **0.0541** |
| Xuño | *0.8967* | *0** | *0.0001** | *0** | - | 0.0052 | 0.0113 | 0 | **0.0187** | 0 | 0.0083 | 0.0008 |
| Perdiguero | *0.0428* | *0** | *0.2627* | *0** | *0.169* | - | 0 | 0 | **0.0350** | 0 | 0.0138 | 0 |
| Arreo | *0.0007** | *0** | *0.004* | *0** | *0.0176* | *0.9911* | - | 0 | **0.0368** | 0 | 0.0131 | 0 |
| Cañas | *0.9629* | *0.3608* | *1* | *1* | *0.9999* | *1* | *1* | - | 0 | 0 | 0 | 0 |
| Mateo | *0** | *0** | *0** | *0** | *0** | *0** | *0** | *0.9615* | - | **0.0304** | **0.0148** | 0.0286 |
| Villar | *0.4047* | *0** | *0.6238* | *0** | *0.8476* | *0.9984* | *0.9565* | *0.6562* | *0.0006** | - | 0 | 0 |
| Valpierre | *0.0001** | *0** | *0** | *0** | *0.0009* | *0.0058* | *0.0107* | *1* | *0** | *0.8821* | - | 0 |
| Valbornedo | *0.2624* | *0** | *0.6843* | *0** | *0.4631* | *0.7494* | *0.7379* | *0.7676* | *0.0016* | *0.5447* | *0.6181* | - |
